# SVIM: Structural Variant Identification using Mapped Long Reads

**DOI:** 10.1101/494096

**Authors:** David Heller, Martin Vingron

**Affiliations:** Computational Molecular Biology Department, Max Planck Institute for Molecular Genetics, Berlin, 14195, Germany

## Abstract

**Motivation:** Structural variants are defined as genomic variants larger than 50bp. They have been shown to affect more bases in any given genome than SNPs or small indels. Additionally, they have great impact on human phenotype and diversity and have been linked to numerous diseases. Due to their size and association with repeats, they are difficult to detect by shotgun sequencing, especially when based on short reads. Long read, single molecule sequencing technologies like those offered by Pacific Biosciences or Oxford Nanopore Technologies produce reads with a length of several thousand base pairs. Despite the higher error rate and sequencing cost, long read sequencing offers many advantages for the detection of structural variants. Yet, available software tools still do not fully exploit the possibilities.

**Results:** We present SVIM, a tool for the sensitive detection and precise characterization of structural variants from long read data. SVIM consists of three components for the collection, clustering and combination of structural variant signatures from read alignments. It discriminates five different variant classes including similar types, such as tandem and interspersed duplications and novel element insertions. SVIM is unique in its capability of extracting both the genomic origin and destination of duplications. It compares favorably with existing tools in evaluations on simulated data and real datasets from PacBio and Nanopore sequencing machines.

**Availability and implementation:** The source code and executables of SVIM are available on Github: github.com/eldariont/svim. SVIM has been implemented in Python 3 and published on bioconda and the Python Package Index.

## 1 Introduction

A typical human genome differs from the reference genome at approximately 4 to 5 million sites amounting to approximately 20 million altered bases [1]. These variations can be categorized into single nucleotide polymorphisms (SNPs), small insertions and deletions (Indels), and structural variations (SVs) affecting a larger number of base pairs. Typically, differences larger than 50bp are considered SVs although definitions vary and sometimes overlap with those of indels.

Studies have shown that in human more base pairs are altered due to structural variation than due to SNPs [2, 3]. Additionally, SVs are enriched 50-fold for expression quantitative trait loci when compared to SNPs [4]. Unsurprisingly, SVs have a major influence on human diversity and are implicated in a wide range of diseases from autism and other neurological diseases to cancer and obesity [5, 3]. Consequently, the characterization of SVs is of major importance to human medicine and genetics alike. It can contribute to the early detection of disorders and can help to elucidate their underlying genetic and molecular processes [6]. In other organisms such as plants, SVs play an equally important role by driving phenotypic variation and adaptation to different environments [7].

Next generation sequencing has enabled the identification of SNPs and small indels to a high resolution. SVs, however, are much harder to detect. One reason is that SVs encompass a diverse range of modifications. While SNPs are simple base pair substitutions, the term “SV” summarizes many different phenomena. Typically, different classes of SVs are distinguished, such as deletions, inversions and insertions. Definitions for some of these classes vary in the literature. For the purpose of this work, we define six different SV classes which are visualized in Figure 1: deletions, cut&paste insertions, tandem and interspersed duplications, inversions and novel element insertions. The main drivers behind interspersed duplications in human are mobile element insertions, such as Alu, LINE1 and SVA elements. They duplicate using retrotransposition and in total represent approximately 25% of all human structural variation [8, 4]. DNA transposons, although now inactive in mammals (excepts bats) are active in plants and lower-order animals [9]. They use a cut&paste mechanism to move in the genome and therefore motivated the inclusion of cut&paste insertions as a separate SV class.

**Figure 1:**
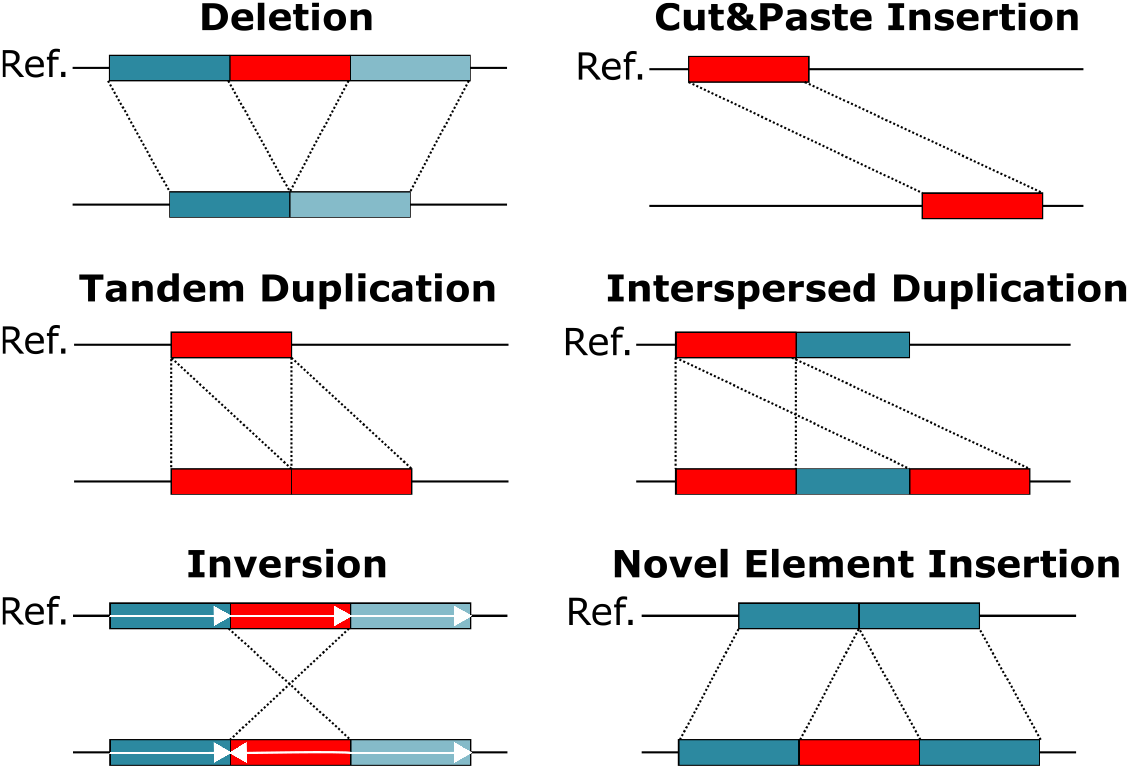
Schematic overview of different SV classes. Structural variations can be categorized into deletions, cut&paste insertions, tandem and interspersed duplications, inversions and novel element insertions. Each SV class is depicted in an individual genome (lower line) when compared to the reference genome (upper line). The region being rearranged is marked in red.

There exists a wide variety of tools for SV calling from short reads [10] but despite ongoing efforts, the discovery of SVs from short-read data remains challenging [11]. Studies have estimated that short-read methods suffer from poor sensitivity down to 10% particularly for small SVs shorter than 1kbp [12, 13]. In contrast to SNPs where discovery and sequence resolution can be performed simultaneously, SVs are discovered mainly indirectly using short paired-end reads. Their alignments are examined for characteristic signatures, such as inconsistently mapping read pairs, split reads and changes in read depth [14]. These signatures can only be indirect evidence in favor of certain SV classes but are unable to fully characterize the SV. The main limitation here is that most SVs are simply larger than the short reads. The accurate detection of SVs is, besides their diversity, hampered by their association with repeat regions, biases in the sequencing technology and the additional complexity of diploidy [15, 16, 17].

To characterize the full spectrum of human genetic variation, long-read sequencing technologies that generate reads with an average length of tens of kilobases show many advantages. The long reads can be mapped with greater accuracy which enables the sequencing of repetitive and low-complexity regions [18, 19]. Unlike with short reads, SVs are often spanned by a single long read. This enables the direct detection and full characterization of the SVs. Consequently, several studies confirmed that a substantial number of SVs that are missed by short-read approaches can be identified with long reads [11, 20, 12]. Two commercial long-read sequencing solutions exist to date: single-molecule real-time (SMRT) sequencing by Pacific Biosciences (PacBio) and nanopore sequencing by Oxford Nanopore Technologies (ONT). Both technologies have the same drawbacks: high error rates of approximately 5-15% with dominating indel errors and still high costs compared to short read sequencing.

Similarly to the detection of SVs from short read data, the first step towards SV detection from long reads is often the alignment of the reads to a reference genome. Depending on the alignment tool used to produce the alignments, SV detection results can vary substantially as Sedlazeck et al. showed for their tool *Sniffles* [21]. In that study, SV-spanning long reads were aligned with seven different aligners. Their results showed that one particular aligner, *NGMLR*, outperformed all the others (including *BWA-MEM*, *Minimap2*, *LAST* and *BLASR*) on the task [21]. In our study, we analyzed read alignments by *NGMLR* to detect SVs. In the Supplementary, however, we include results for *Minimap2* which is an order of magnitude faster than *NGMLR* [22].

Read alignments alone are not sufficient to detect and characterize SVs. Dedicated SV callers are needed to collect and interpret evidence from the read alignments. Recently, three methods have been developed for calling SVs based on long reads [23]. *PBHoney* and *SMRT-SV* are designed specifically for PacBio reads while *Sniffles* supports PacBio and ONT reads [24, 12, 21].

*PBHoney* comprises two different variant identification approaches [24]. The first approach, *PBHoney-Spots*, exploits the stochastic nature of the errors in PacBio reads. It scans read alignments (usually produced by the read aligner *BLASR*) and recognizes SVs by an increase in error and a subsequent decrease in error along the reference sequence. The second approach, *PBHoney-Tails*, analyzes the soft-clipped (i.e. unmapped) read tails from a *BLASR* alignment. It extracts such tails from the BLASR output and realigns them to the reference. Then, SVs are detected by clustering the resulting piece-alignments based on their location and orientation.

*SMRT-SV* scans PacBio alignments for SV signatures, such as spanned deletions, spanned insertions and soft-clipped read tails [12]. Clusters of such events are validated with a local de-novo assembly of the reads overlapping the locus and subsequent alignment of the assembly to the reference.

*Sniffles* uses signatures from split-read alignments, high-mismatch regions, and coverage analysis to identify SVs [21]. To overcome the high error rate in the reads, it evaluates candidate SVs based on features such as their size, position and breakpoint consistency.

All three methods regard structural variation (i.e. deletions, insertions, inversions) as rearrangements occurring in a single genomic locus. However, structural variation often involves multiple genomic loci, such as for a mobile element which is reverse-transcribed from a source region and inserted at another location. The higher read lengths of PacBio and ONT reads allow to link both loci much more efficiently and confidently than was possible with short paired-end reads. Nevertheless, existing methods ignore this type of information and are only able to detect the isolated destination location of the mobile element insertion.

In this study, we introduce SVIM, a computational method for the sensitive detection and accurate classification of five different classes of SVs from long read sequencing data. We describe the three core components of the approach and our methodology for evaluation on simulated and real datasets. Our results demonstrate that SVIM reaches substantially higher recall and precision than existing tools for SV detection from long reads. Unlike other methods, SVIM has been specifically designed to distinguish three separate classes of large insertions: interspersed duplications, tandem duplications and insertions of novel elements. To our knowledge, it is the only tool capable of identifying not only the insertion location of an interspersed duplication but also its potential genomic origin. We demonstrate this capability on a small number of high-scoring interspersed duplications identified in the NA12878 individual. Furthermore, we compare SV callsets produced by SVIM on reads from PacBio and Nanopore data. Finally, we compare the runtimes of different SV callers including SVIM.

## 2 Methods

SVIM implements a pipeline of three consecutive components (see Fig. 2). First, SV signatures are collected from each individual read in the input SAM/BAM file (COLLECT). Secondly, the detected signatures are clustered using a graph-based clustering approach and a novel distance metric for SV signatures (CLUSTER). Thirdly and lastly, multiple SV events are merged and classified into higher-order events (i.e. events involving multiple regions in the genome) such as duplications (COMBINE). The three components are explained in the following.

**Figure 2:**
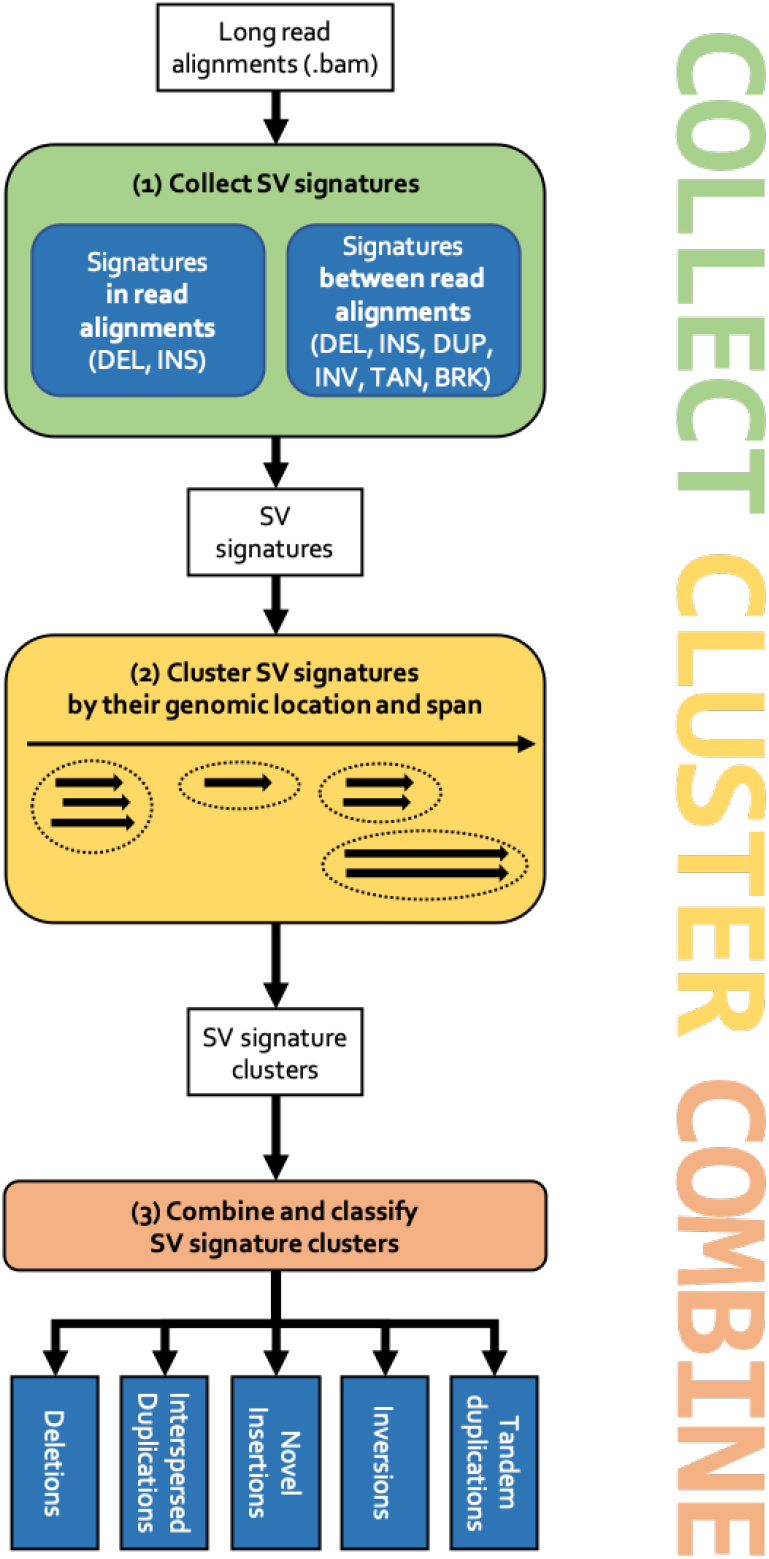
The SVIM workflow. (1) Signatures for SVs are collected from the input read alignments. SVIM collects them from within alignments (intra-alignment signatures) and between alignments (inter-alignment signatures). (2) Collected signatures are clustered based on their genomic position and span. (3) Signature clusters from different parts of the genome are combined to distinguish five different classes of SVs: deletions, interspersed duplications, novel insertions, inversions and tandem duplications.

### 2.1 Collection of SV signatures from individual reads

SVIM analyzes read alignments in SAM/BAM format [25] from a read aligner. Modern aligners, such as *NGMLR* and minimap2, try to find good linear alignments of entire reads. Nevertheless, they will split a chimeric read if its different segments can be better aligned separately. Due to these split alignments, the SAM/BAM output from these aligners can contain multiple alignments for each read (one for each aligned read segment). SVIM extracts signatures for SVs from the SAM/BAM file by analyzing one read at a time. We define SV signatures as discordant alignments of a read that point to the presence of one or several possible SVs in the sequenced genome. SVIM searches for two types of signatures:

**Intra-alignment signatures** are large alignment gaps in the reference or in the read. They can be found in the CIGAR strings of individual SAM/BAM entries.
**Inter-alignment signatures** are discordant relative alignment positions and orientations of a read’s alignment segments. To illustrate this type of evidence, imagine an inversion that is spanned by a single read. The aligner will split the read into three alignment segments: one segment upstream of the inversion, another segment for the inverted region, and a third segment downstream of the inversion. Due to the inversion, the middle segment will have a different mapping orientation than the other two pieces. This and other types of inter-alignment signatures are detected by SVIM in a heuristic fashion.

This analysis yields 6 different types of SV signatures: (1) deleted regions (DEL), (2) inserted regions (INS), (3) inserted regions with detected region of origin (DUP), (4) inverted regions (INV), (5) tandem duplicated regions (TAN) and (6) translocation breakpoints (BRK). Some of these evidence types (e.g. inverted regions) indicate one particular SV class. Others could indicate several possible SV classes. An inserted region, for instance, can indicate both a duplication or a novel element insertion.

### 2.2 Clustering of SV signatures

The collection of signatures from the alignments is only the first step to accurately detect SVs. Subsequently, signatures from multiple reads need to be merged and criteria have to be found to distinguish correct signatures from multiple types of error artifacts (e.g. sequencing error, alignment error). To achieve this, we combine a graph-based clustering approach with a novel distance metric for SV signatures. The aim is to merge signatures of the same SV even if their positions vary slightly due to sequencing or alignment errors. At the same time, signatures from separate SVs need to be kept separate even if the two SVs lie close to each other.

The collected SV signatures can be viewed as quadruples *S_i_* = (*T_i_,C_i_, B_i_, E_i_*) where *T* is one of the six different signature types defined above, *C* is the chromosome and *B* and *E* are the genomic start (begin) and end positions. One of the few distance metrics defined for such genomic intervals is the Gowda-Diday distance [26]. It combines (a) the distance between two intervals, (b) their span difference, and (c) their degree of overlap into a single numeric distance value. In our type of data (i.e. long read alignments), however, we often observe little to no overlap between signatures originating from the same SV but from different long reads (see Suppl. Fig. S1). Nevertheless, signatures from the same SV often possess similar positions and spans.

Therefore, we introduce *span-position distance* as a novel distance metric for SV signatures. For two SV signatures *S*_1_ and *S*_2_, the span-position distance *SPD* consists of two components *SD* and *PD*: 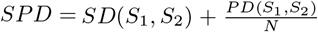. *SD* is the difference in span between both signatures (normalized to [0,1)) and is defined as 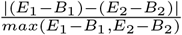. *PD* is the difference in position between both signatures and is defined as 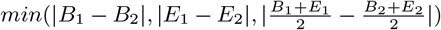. *N* is a user-defined normalization constant which regulates the relative importance of *SD* and *PD*. In our analyses, setting *N* = 900 returned the best results. Intuitively, this setting means that two signatures that are 900bp apart (*PD* = 900) but have the same span (*SD* = 0) would have the same SPD as two signatures with extremely different spans (*SD* ≈ 1) but the same position (*PD* = 0).

To perform clustering, we follow a graph-based approach similar to the one used by the variant finder *CLEVER* [27]. Initially, we transform the set of collected SV signatures into an undirected graph. While *CLEVER* identifies nodes with alignments of short paired-end reads, each node in our graph represents an SV signature. We draw an edge between two nodes (i.e. signatures) if the span-position distance between the two signatures is smaller than a user-defined threshold *T*. Systematic evaluation of different settings for this parameter yielded *T* = 0.7 as an optimal setting for our human datasets (data not shown). An edge between two nodes expresses our confidence that the two signatures represented by the nodes express the same SV allele. From the graph, we produce signature clusters by extracting maximal cliques with an efficient implementation of the Bron-Kerbosch algorithm [28, 29]. As a consequence, each signature cluster is a maximal group of SV signatures that can be jointly assumed to express the same SV in the donor genome.

Finally, SVIM computes a score for each cluster based on four features:

1. The number *n* ∈ (0, 40] of signatures in the cluster where at most 20 of each class (intra-alignment or inter-alignment) are taken into account.
2. An additional bonus *b* ∈ [0, 30] for the existence of at least one signature from each of the two classes. One or more intra-alignment signatures earn a bonus of 10 while one or more inter-alignment signatures earn an additional bonus of 20.
3. A score *s_p_* ∈ [0, 10] based on the standard deviation *s_pos_* of the genomic positions of the signatures in the cluster normalized by their average span.

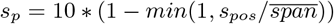
4. A score *s_s_* ∈ [0, 20] based on the standard deviation *s_span_* of the genomic spans of the signatures in the cluster normalized by their average span.

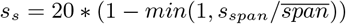 By summing up these four components we obtain a score *S G* (0, 100] to discern trustworthy signature clusters from artifacts, such as sequencing or alignment artifacts. Trustworthy events are characterized by many intra- and inter-alignment signatures that exhibit high concordance regarding their genomic position and span.

### 2.3 Combination and classification of SVs into five SV classes

The third component in the workflow analyzes and combines the SV signature clusters to classify events into five SV classes: deletions, inversions, novel element insertions, tandem duplicaitons and interspersed duplications. Because the confident distinction of interspersed duplications and cut&paste insertions solely based on sequencing reads is impossible, we classify both as interspersed duplications. Nevertheless, we annotate duplications where the region of origin seems to be deleted in the sequenced individual (i.e. a deletion overlaps the genomic origin) as potential cut&paste insertions. While inversions, deletions and tandem duplication signature clusters can be directly reported as inversions, deletions and tandem duplications, respectively, the other three signature classes (INS, DUP and BRK) are more complex. The reason is that interspersed duplications are not characterized by only one genomic region but two – a genomic origin and a genomic destination. To capture and classify these higher-order events, SVIM needs to combine multiple signature clusters and therefore makes the following distinctions (see also Fig. 3):

- Insertion signature clusters with detected region of origin (DUP) are called as interspersed duplications. If the genomic origin overlaps a deletion call, the duplication is marked as potential cut&paste insertion.
- Inserted region signature clusters (INS) that are close to matching translocation breakpoints (BRK) are called as interspersed duplications. If the genomic origin (as defined by the translocation breakpoints) overlaps a deletion call, the duplication is marked as potential cut&paste insertion.
- The remaining inserted region signature clusters (INS) are called as novel element insertions.

**Figure 3:**
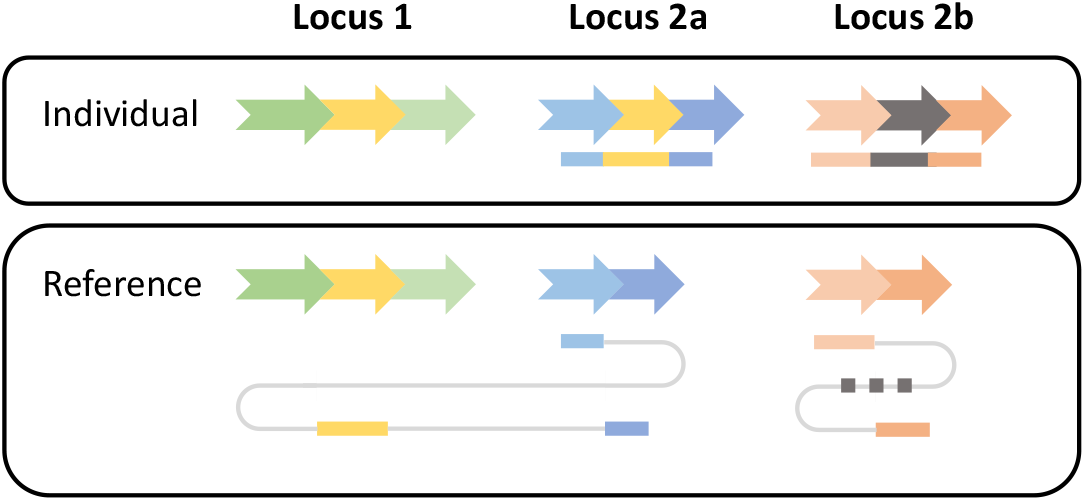
Read signatures for an interspersed duplication and a novel element insertion. A genomic segment (yellow arrow) has been copied from locus 1 to locus 2a in an individual genome. Additionally, a novel genomic segment (gray arrow) has been inserted in locus 2b. Two reads are generated from the individual (top) and mapped to the reference genome (bottom). The first read (blue-yellow) consists of three segments. They are mapped individually to the reference genome. The two blue segments are mapped to locus 2a exhibiting an insertion signature. The yellow segment is mapped to locus 1 indicating the origin of the insertion. The second read (orange-gray) exhibits a similar insertion signature at locus 2b but as the inserted gray segment is unmapped its origin cannot be determined.

### 2.4 Implementation and usage

SVIM has been implemented in Python and is available at github.com/eldariont/svim. It can be easily installed via bioconda or the Python Package Index (PyPI). As input, SVIM expects either raw reads (in FASTA or FASTQ format) and a reference genome (in FASTA format) or already aligned reads in BAM format. It outputs detected SVs in five separate BED files (one for deletions, interspersed and tandem duplications, inversions and novel insertions, respectively). Additionally, a VCF file with all SV results is produced.

### 2.5 Evaluation methodology

In this study, we compared our tool, SVIM (v0.4.1), to three other SV detection methods: *PBHoney-Spots, PBHoney-Tails* (both PBSuite v15.8.24) and *Sniffles* (v1.0.8). All three tools are designed for the application on long read sequencing data. For *Sniffles* and SVIM, reads were aligned with *NGMLR* (v0.2.7) or *minimap2* (v2.12-r836-dirty). For *PBHoney*, reads were aligned with *BLASR* (v5.3.4323a52). We did not compare against short read SV callers because they have been shown to exhibit lower recall than methods relying on long reads [13, 12, 21]. We also did not compare against *SMRT-SV* because it is not a stand-alone tool but a software pipeline applying several alignment, detection, and assembly steps with various other tools. It detects only three SV classes and is computationally more demanding than pure alignment-based tools.

We evaluated all tools on two types of data. Firstly, we generated a simulated genome from which we sampled in-silico PacBio sequencing reads with known SVs. This provided us with a complete set of fully characterized SVs for evaluation. Secondly, we used publicly available sequencing reads from PacBio and Nanopore sequencers. We compared the precision and recall of the three methods. Precision is defined as the fraction of detected SVs that are correct (requiring 50% reciprocal overlap between detected and correct SVs). Recall is defined as the fraction of correct SVs that have been detected (with 50% reciprocal overlap). Results for a more lenient and a more stringent overlap requirement of 1% and 90%, respectively, can be found in the Supplementary Material. Both precision and recall require a suitable gold standard set of high-confidence SVs for the given genome (i.e. a set of correct SVs).

As expected, recall and precision reached by the different tools depend heavily on tool parameters, particularly score or support thresholds. More relaxed thresholds (i.e. yielding more SVs) increase recall but decrease precision while stricter cutoffs achieve the opposite. Consequently, we ran all four tools with different settings of their most important parameter: For SVIM we applied different score cutoffs (0 to 100). *Sniffles* was run with different settings of the *min_support* parameter (1 to 60). For *PBHoney-Spots*, we varied the *minErrReads* parameter and for *PBHoney-Tails* we varied the *minBreads* parameter (both 1 to 60). We visualized the performance of the tools by plotting each parameter setting as a distinct point in Figures 4–6. Besides that one parameter, we used the default settings for all other tool parameters except PB-Honey Spots’ *spanMax* parameter which we set to 100,000 (100kb).

**Figure 4:**
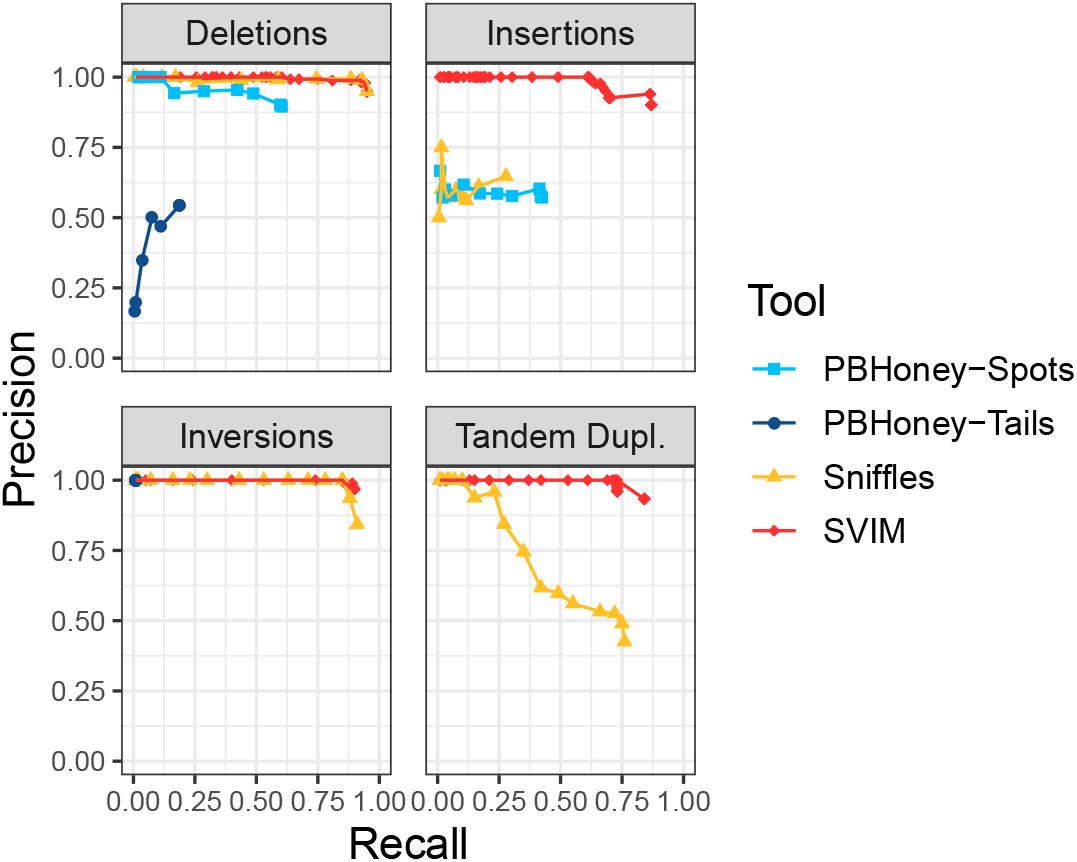
Comparison of SV detection performance on a 6x coverage homozygous simulated dataset. SVIM consistently yielded better recall (x-axis) and precision (y-axis) than the other tools for the recovery of inserted regions and tandem duplications. For the recovery of deletions and inversions, *Sniffles* reached the same recall as SVIM. The different points for each tool represent multiple settings of the tools’ most important parameters (see Sec. 2.5). *PBHoney-Spots* only detects deletions and inserted regions and *PBHoney-Tails* does not detect duplications. Recall and precision were calculated using a required reciprocal overlap of 50% between variant calls and the original simulated variants.

#### 2.5.1 Simulated data

We simulated 600 homozygous SVs by altering the sequence of chromosomes 21 and 22 in the hg19 reference genome. More precisely, we implanted 200 deletions, 100 inversions, 100 tandem duplications, and 200 interspersed duplications with the R package *RSVSim* [30]. The package estimates the distribution of SV sizes from real datasets and simulates the association of SVs to various kinds of repeats. The resulting genome contained SVs between 50bp and 10kbp in size. Subsequently, reads were simulated from this genome to generate 10 different datasets with coverages between 6x and 60x with the tool *SimLoRD* [31]. *SimLoRD* imitates the error model of SMRT reads to simulate realistic PacBio reads.

To simulate heterozygous SVs, we adapted the previously described approach only slightly. Instead of sampling all reads from the altered reference genome, half of the reads were sampled from the original refer ence genome. Consequently, reads from the original (wildtype) reference genome and the altered genome each amounted to 50% of the total coverage.

The comparison between different tools was complicated by the fact that each tool is designed to detect different SV classes. *PBHoney* is able to detect deletions, inserted regions, inversions and translocation breakpoints. *Sniffles* is additionally capable of identifying tandem duplications and complex events. Because only SVIM distinguishes between duplications and novel element insertions, we compared the tools on four common basic SV classes in the simulated datasets: deletions, inserted regions (i.e. inserted sequence from duplications and novel element insertions), inversions and tandem duplications. Because *Sniffles* tends to call intra-chromosomal duplications as very large deletions or inversions (see github.com/fritzsedlazeck/Sniffles/issues/23), we omitted deletion and inversion calls by *Sniffles* that were larger than 100kbp to ensure a fair comparison. To obtain calls of inserted regions from SVIM, we use the union of its interspersed duplication and novel element insertion calls.

#### 2.5.2 Real data

Simulation cannot reflect all aspects of biological data. Therefore, we used real PacBio and Nanopore data for the second part of our analysis. This part consisted of three separate experiments. For the first two, we utilized a real 53x coverage dataset of the NA12878 individual from a PacBio RS II machine (Genome in a Bottle consortium; Accession SRR3197748) [32]. To assess the influence of sequencing coverage on SV detection performance, we produced a corresponding low-coverage subset of the dataset by sampling reads randomly to 6x coverage. With these two PacBio datasets, we performed two separate analyses. Firstly, we evaluated our method with a published benchmark sample of 2676 high-confidence deletions and 68 high-confidence insertions [33]. Secondly, we implanted SVs into the reference genome and aligned the PacBio reads to this altered reference. Implanting an SV into the reference genome causes the original reads to contain the inverse of the SV that was implanted. With this approach, three types of SVs were simulated: 1) 200 deletions were simulated by inserting sequence into the reference genome. 2) 100 inversions were simulated by inverting regions in the reference. 3) 200 insertions were simulated by deleting regions in the reference. Unfortunately, duplications could not be simulated because this would have required the identification and alteration of existing duplications in the reference genome.

In a third experiment, we compared the 53x coverage PacBio dataset of the NA12878 individual with a 26x coverage Nanopore dataset of the same individual ([34], release 5). We evaluated our method with the high-confidence callset described above and analyzed the overlap between the three callsets (PacBio, Nanopore and high-confidence callset).

The NA12878 datasets are more realistic than the simulated dataset but impose the limitation that there exists no complete gold standard set of SVs. As a consequence of using an incomplete gold standard for evaluation, precision could not be accurately measured. Putative “false positives” could have been true but simply not contained in the incomplete gold standard. Therefore, we compared the tools only based on their recall in relation to the number of calls.

## 3 Results

### 3.1 Evaluation with simulated reads

As described in the Methods section, we implanted SVs from four different classes into a reference genome. Reads sampled from this synthetic genome were then analyzed with SVIM, *PBHoney-Tails*, *PBHoney-Spots* and *Sniffles*. Results for the 6x coverage homozygous dataset can be found in Figure 4. For a comparison of results across all coverages from 6x to 60x see Suppl. Fig. S2.

Regardless of coverage, SVIM achieved substantially better results than all other tools in the recovery of inserted regions and tandem duplications. With 6x coverage and homozygous SVs, SVIM reached average precisions (AP) of 86% (inserted regions), and 83% (tandem duplications) for the two classes while the second best tools, *PBHoney-Spots* and *Sniffles* respectively, reached 25% and 54%. In the recovery of deletions and inversions, SVIM and *Sniffles* reached equal results with average precisions of 94% (deletions) and 90% (inversions), respectively. In our experiments, *PBHoney-Tails* performed very poorly across all settings. It did detect only very few inserted regions, suffered from very low recall for inversions and poor precision for deletions. All these trends remain true for higher coverages as well (see Suppl. Fig. S2).

The simulated heterozygous dataset yielded similar results to those of the homozygous dataset (see Suppl. Fig. S5). While all tools reached slightly lower precision and recall, SVIM still outperformed the others for inserted regions (AP=68% for 6x coverage) and tandem duplications (AP=76%). In the detection of deletions and inversions, however, *Sniffles* and SVIM reached nearly equal results (AP=90% and AP=87%, respectively).

We explored whether more lenient (1%) or stringent (90%) overlap requirements for the calls would change the results (see Suppl. Fig. S3, S4, S6, and S7). As it turned out, the overlap requirement had little effect on *Sniffles* and SVIM. Only *PBHoney-Spots* produced substantially worse results for more stringent overlap requirements suggesting that the tool has trouble finding accurate SV breakpoints.

To measure the influence of the input read alignments on SV calling, we also compared results for two long read aligners, *NGMLR* and *minimap2* (see Suppl. Fig. S8 and S9). The results indicate that SVIM is relatively robust to the choice of the aligner but benefits slightly from the more accurate alignment of reads covering insertions and tandem duplications by *NGMLR. Sniffles*, however, reaches considerably higher recall for insertions when analyzing alignments by *minimap2* compared to *NGMLR*. Visual inspection of the alignments revealed a difference in the way that reads covering insertions are aligned. While *minimap2* expresses insertions mainly as long reference gaps in the CIGAR string, *NGMLR* tends to split reads at insertions. Because *Sniffles* does not call insertions of sequence existing somewhere else in the genome (i.e. interspersed duplications) from split alignments, it reaches higher recall with *minimap2*.

### 3.2 Evaluation with real reads and high-confidence calls

While simulated datasets enable the comprehensive comparison of tools in a controlled and precise manner, they cannot reflect the full complexity of real sequencing data. Therefore, we analyzed a publicly available 53x coverage dataset of a human individual from a PacBio RS II machine and a random 6x coverage subset (see Methods section). To evaluate the detection performance of our tool, we first used a published benchmark set of 2676 high-confidence deletions and 68 high-confidence insertions.

Among all tools, SVIM was the most consistent across the different settings (see Fig. 5). It recovered substantially more deletions from the high-confidence call set than the other tools with the same number of SV calls. To reach a recall of more than 50%, SVIM needed 1932 / 2577 calls (53x/6x coverage) while *Sniffles* needed 4320 / 6333 calls. *PBHoney-Spots* needed even 5062 calls (53x coverage) and *PBHoney-Tails* did not reach this level of recall at all. A recent study by the Human Genome Structural Variation Consortium (HGSVC) used a multi-platform de-novo assembly approach for SV detection and found an average of 12,680 deletions per individual [35]. When we select tool thresholds closest to this mark, SVIM, *Sniffles, PBHoney-Spots* and *PBHoney-Tails* recover 97%, 97%, 80% and 46% of the high-confidence deletions from the full coverage dataset, respectively. All tools miss high-confidence calls across the entire size range (50bp – 140kb). But while the false negatives of the first three tools are evenly distributed across the size spectrum, *PBHoney-Tails* particularly misses small events. For instance, it misses all high-confidence calls smaller than 100bp and 69% of calls between 100bp and 500bp but only 24% of calls between 500bp and 1kb.

**Figure 5:**
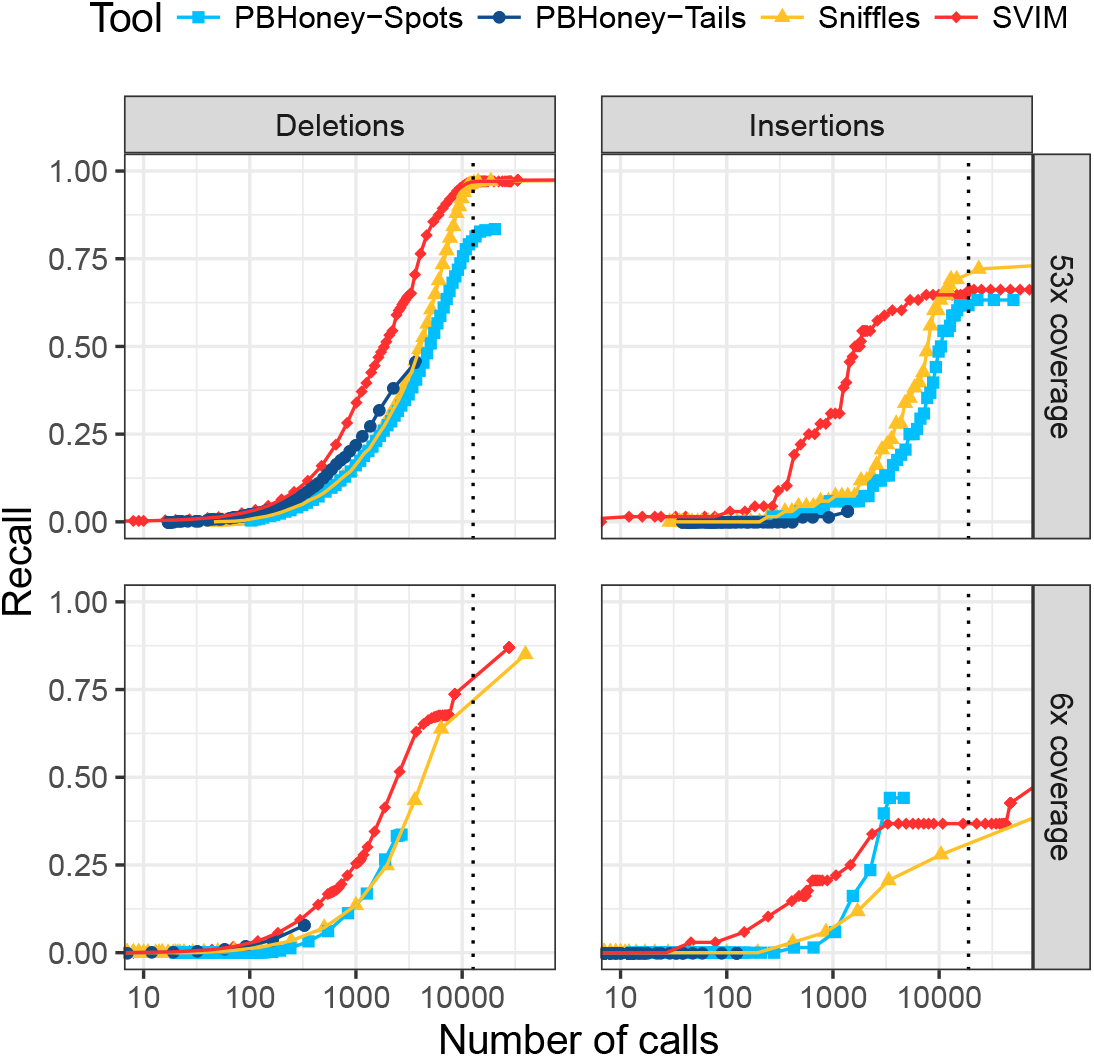
Comparison of recall on a 53x coverage public PacBio dataset and a 6x coverage subset with 2676 high-confidence deletion and 68 insertion calls. For each tool and different thresholds, the number of SV calls with score above the threshold (log-scale) is plotted against the recall. The upper and lower panels show performance on the full dataset and a randomly sampled 6x coverage subset of the data, respectively. SVIM reached the same recall with fewer calls than other tools. The vertical dotted lines denote the average number of deletions and insertions to expect in an individual as recently reported using a de-novo assembly approach [35]. Recall was calculated using a required reciprocal overlap of 50% (deletion calls) and 1% (insertion calls), respectively, between variant calls and the gold standard variants.

Although the results for insertions need to be considered with greater caution due to the small size (n=68) of the high-confidence call set, SVIM reached a higher recall than all other tools for small numbers of calls. When we again select tool thresholds closest to the estimate of 18,919 insertions per individual from the HGSVC study [35], SVIM, *Sniffles, PBHoney-Spots* and *PBHoney-Tails* recover 66%, 72%, 62% and 3% of high-confidence insertions from the full coverage dataset, respectively. Again, all tools miss high-confidence calls across the entire size range of the callset (12bp – 379bp).

### 3.3 Evaluation with real reads and an altered reference genome

As described in the Methods section, we obtained another reliable gold standard set of SVs (deletions, inversions, insertions) by implanting SVs into the reference genome and aligning the PacBio reads (53x and 6x coverage) to this altered reference. We evaluated all combinations of the three SV types and the two coverages. SVIM outperformed the other tools (see Fig. 6) in all six of these settings. In the recovery of deletions and inversions, SVIM reached a substantially higher recall than *PBHoney*. It also needed fewer SV calls to reach similar recall than *Sniffles* while the difference decreased for higher recall. The most striking difference was observed for the detection of insertions. While SVIM reached a recall of 84% and 43% with 20,000 calls (53x and 6x coverage, respectively), *PBHoney-Spots* reached 61% and 25% and *Sniffles* detected only 57% and 29% with the same number of calls. For full coverage, SVIM needed 2480 calls to reach a recall of 50% while *Sniffles* and *PBHoney-Spots* needed both more than 10,000 calls.

**Figure 6:**
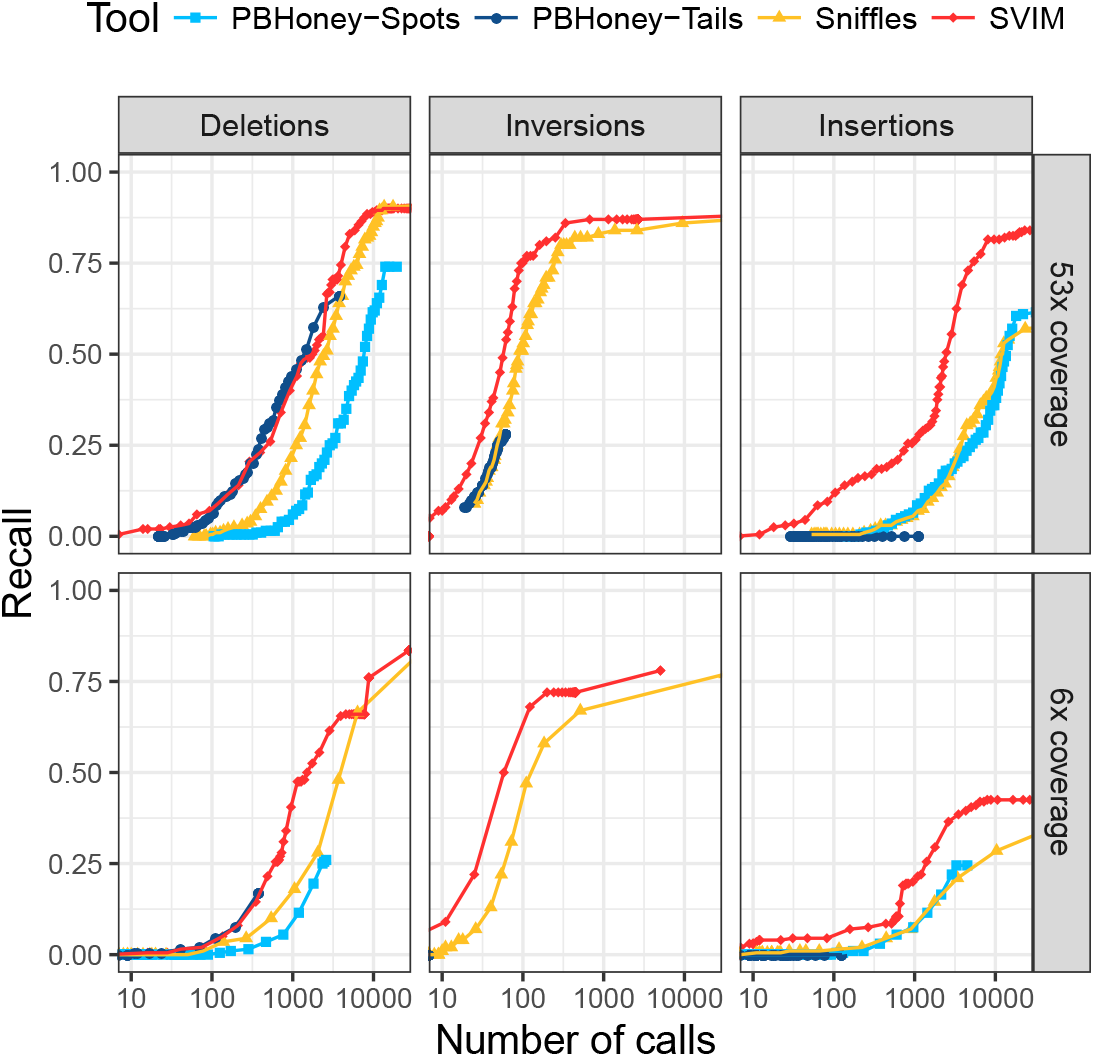
Comparison of recall from NA12878 reads aligned to an altered reference genome. For each tool and different thresholds, the number of SV calls with score above the threshold (log-scale) is plotted against the recall. The upper and lower panels show performance on the full dataset and a randomly sampled 6x coverage subset of the data, respectively. In all six panels, SVIM outperformed all the other tools and reached substantially higher recall for similar numbers of calls. The improvement was most prominent for insertions. Recall was calculated using a required reciprocal overlap of 50% between variant calls and the original implanted variants.

### 3.4 Interspersed duplications in NA12878

SVIM’s ability to link the genomic origin and destination of an interspersed duplication can yield interesting insights into the dynamics of genomic rearrangements. Our analysis of the NA12878 PacBio dataset with SVIM identified 27 high-confidence interspersed duplications with a score > 30 (Supplementary Table 1). The genomic origin of 19 of them overlapped annotated retrotransposons. Among those, 10 and 2 represented complete and incomplete Alu insertions, respectively. 2 and 2 represented insertions of complete and incomplete LINE1 elements, respectively. 2 represented complete SVA elements and another one represented human endogenous retrovirus HERVK14. Strikingly, six duplications occurred from regions of the genome without annotated repeat elements indicating other formation mechanisms. Finally, we observed two duplications in the untranslated regions of three genes, BAZ2A, RBMS2 and PCMTD1.

### 3.5 Comparison of PacBio and Nanopore sequencing data

SVIM can detect SVs from both PacBio and Nanopore data. An evaluation with real reads and high-confidence calls demonstrated that SVIM’s performance on a 26x coverage Nanopore dataset is comparable to its performance on the 53x coverage PacBio dataset (see Suppl. Fig. S14). When we compared both SVIM callsets with the high-confidence callset, we found that all three callsets together yielded a total of 45,729 SVs (score cutoff of 40; see Figure 7). 22,461 or 49% of the calls were unique to one of the callsets with 13,385 and 9,017 SVs detected exclusively from the PacBio and Nanopore reads, respectively. However, 23,248 or 51% of the calls were made on both PacBio and Nanopore reads. It is reassuring that the vast majority (97%) of high-confidence calls were detected by both sequencing technologies.

**Figure 7:**
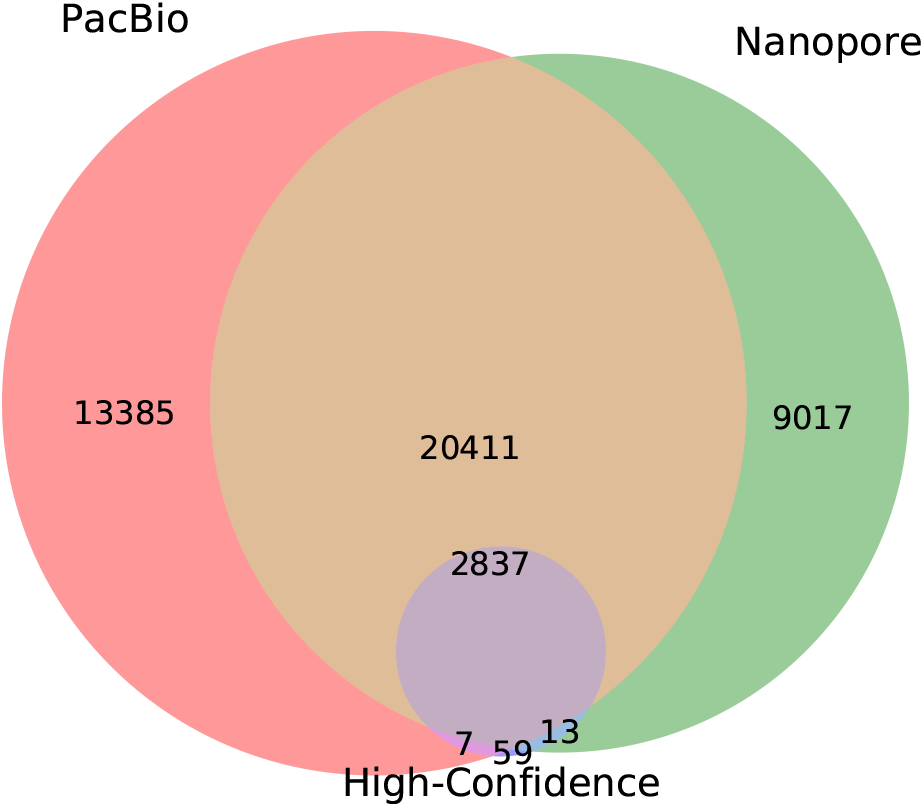
Venn diagram of three SV callsets for NA12878: SVIM calls on a 53x coverage PacBio dataset, SVIM calls on a 26x coverage Nanopore dataset and high-confidence calls from [33]. Callsets were produced by merging SVIM calls with a score ≥ 40 for deletions, interspersed duplications and novel element insertions. Subsequently, the diagram was generated using *pybedtools* [36] and *matplotlib_venn.*

PacBio and Nanopore sequencing exhibit similar error rates but slightly different error modes. While PacBio produces more insertion than deletion errors [37], Nanopore shows the opposite tendency [38]. In concordance to these biases, SVIM detected 17,292 deletions from the Nanopore reads but only 12,782 deletions from the PacBio data (see Suppl. Fig. S15). Conversely, it detected 23,858 insertions from PacBio but only 14,986 insertions from Nanopore data (see Suppl. Fig. S16). Consequently, the majority of PacBio-only calls were insertions (90%) and the majority of Nanopore-only calls were deletions (65%). We could confirm the finding by Sedlazeck et al. that a large fraction (80%) of Nanopore-only calls lay in simple tandem repeats in contrast to only 35% of Pacbio-only calls.

### 3.6 Runtime evaluation

We compared the runtimes of *PBHoney-Spots*, *PBHoney-Tails*, *Sniffles*, and SVIM on the same NA12878 dataset (53x coverage). *Sniffles* and SVIM were given input alignments produced by *NGMLR* while *PBHoney-Spots* and *PBHoney-Tails* were given *BLASR* alignments. The runtime was measured on an AMD EPYC 7601 (128 cores, 2.7 GHz, 1 TB memory). Only the runtime of SV detection was measured, excluding the time required for producing the alignments. All four tools analyzed the entire dataset in under 3 hours (see Tab. 1). *PBHoney-Tails*, *Sniffles*, and SVIM use only a single core and took 57, 160, and 156 minutes, respectively. *PBHoney-Spots* is the only tool benefiting from multiple cores and took 145 minutes on 4 cores (608 min on only 1 core).

**Table 1:** Runtime comparison on the 53x coverage NA12878 PacBio dataset. Only the runtime of each tool is measured excluding the prior read alignment.

| Tool | Threads | CPU time (min) | Wall clock time (min) |
| --- | --- | --- | --- |
| PBHoney-Spots | 1 | 601 | 608 |
| PBHoney-Spots | 2 | 561 | 284 |
| PBHoney-Spots | 4 | 558 | 145 |
| PBHoney-Tails | 1 | 56 | 57 |
| Sniffles | 1 | 159 | 160 |
| SVIM | 1 | 155 | 156 |

## 4 Discussion

Structural variation is, besides single-nucleotide variation and small indels, one of the main classes of genetic variation. The influence of SVs on human phenotype and disease makes them an important research target but their unique properties complicate their detection and characterization. Particularly SV detection methods using short read technology suffer from low sensitivity. Long read sequencing technologies such as PacBio SMRT sequencing and ONT Nanopore sequencing have the potential to alleviate these problems. In this study, we introduced the novel SV detection method SVIM. It employs a three-step pipeline to collect, cluster and combine SV signatures from long reads.

A comparison of SVIM with three competing tools on simulated and real data demonstrated that our method combines high sensitivity with high precision. Across all tools, deletions were the easiest to detect. Consequently, Sniffles and SVIM reached almost perfect precision and recall on the simulated data. On the real datasets, both tools still reached a recall of over 90% when setting thresholds using the HGSVC estimate of 12,680 deletions per individual [35]. This level of recall was maintained regardless of sequencing technology and evaluation method (high-confidence callset or altered reference). SVIM generally required fewer calls to reach the same recall as the other tools indicating that the best-scoring SVIM calls are more enriched in true variants than the other tools’ callsets of similar size. For the identification of inversions, *Sniffles* and SVIM exhibited equally strong performance although SVIM showed a slightly higher recall in the evaluation with an altered reference. It needs to be noted, however, that the evaluation of inversions had to rely fully on simulation due to the lack of a suitable gold standard set.

Differences between SVIM and the other tools were most prominent for insertions (i.e. interspersed duplications and novel element insertions). Across all simulations and real data evaluations, SVIM outperformed the other tools by a wide margin. The difference to *Sniffles* can be largely explained by their approach of analyzing split alignments. From such alignments, *Sniffles* only calls insertions of novel elements but does not detect insertions of sequence existing somewhere else in the genome (e.g. from mobile elements). The detection performance of tandem duplications could only be evaluated in the simulated dataset again due to the lack of a gold standard. What we observed is a big difference in precision between SVIM and *Sniffles* due to a large number of erroneous duplication calls by Sniffles.

What sets SVIM apart from existing SV callers is not only its improved detection performance but also its ability to distinguish three different classes of insertions purely based on alignment information. SVIM enables researchers to determine whether an insertion event is due to a tandem duplication, an interspersed duplication or the insertion of a novel element. Moreover, SVIM identifies the genomic origin of duplications which facilitates their functional annotation, e.g. into different classes of mobile elements.

Because SVIM, similar to other SV callers, analyzes read alignments it depends on the correctness of these alignments and inherits the limitations of the used read alignment method. One of these limitations originates from the repetitive nature of many genomes which keeps repetitive read segments from being mapped confidently. This can affect SVIM’s sensitivity but might also cause SVIM to classify an interspersed duplication as a novel insertion if the inserted segment cannot be uniquely mapped. This might particularly affect mobile element insertions whose individual copies are highly similar. Currently, SVIM is unable to detect chromosomal translocations and nested structural variants. We intend to add this functionality in the future. Additionally, we plan to implement genotyping capabilities for the detected variants in an upcoming release of SVIM.

## Supporting information

## Acknowledgements

The authors wish to thank Giuseppe Gallone, M-Hossein Moeinzadeh, Rosario Piro, and Knut Reinert for helpful discussions.

## References

[1] 1000 Genomes Project Consortium. A global reference for human genetic variation. Nature, 526(7571):68–74, 2015.

[2] Richard Redon, Shumpei Ishikawa, Karen R Fitch, Lars Feuk, George H Perry, T Daniel Andrews, Heike Fiegler, Michael H Shapero, Andrew R Carson, Wenwei Chen, et al. Global variation in copy number in the human genome. Nature, 444(7118):444–454, 2006.

[3] Joachim Weischenfeldt, Orsolya Symmons, Francois Spitz, and Jan O Korbel. Phenotypic impact of genomic structural variation: insights from and for human disease. Nat. Rev. Genet., 14(2):125–138, 2013.

[4] Peter H Sudmant, Tobias Rausch, Eugene J Gardner, Robert E Handsaker, Alexej Abyzov, John Huddleston, Yan Zhang, Kai Ye, Goo Jun, Markus Hsi-Yang Fritz, et al. An integrated map of structural variation in 2,504 human genomes. Nature, 526(7571):75–81, 2015.

[5] Jonathan Sebat, B Lakshmi, Dheeraj Malhotra, Jennifer Troge, Christa Lese-Martin, Tom Walsh, Boris Yamrom, Seungtai Yoon, Alex Krasnitz, Jude Kendall, et al. Strong association of de novo copy number mutations with autism. Science, 316(5823):445–449, 2007.

[6] Manuel L Gonzalez-Garay. The road from next-generation sequencing to personalized medicine. Pers. Med., 11(5):523–544, 2014.

[7] Rachit K Saxena, David Edwards, and Rajeev K Varshney. Structural variations in plant genomes. Brief. Funct. Genomics, 13(4):296–307, 2014.

[8] Chip Stewart, Deniz Kural, Michael P Strömberg, Jerilyn A Walker, Miriam K Konkel, Adrian M Stütz, Alexander E Urban, Fabian Grubert, Hugo YK Lam, Wan-Ping Lee, et al. A comprehensive map of mobile element insertion polymorphisms in humans. PLoS genetics, 7(8):e1002236, 2011.

[9] Cheng Ran Lisa Huang, Kathleen H Burns, and Jef D Boeke. Active transposition in genomes. Annual review of genetics, 46:651–675, 2012.

[10] Stephan Pabinger, Andreas Dander, Maria Fischer, Rene Snajder, Michael Sperk, Mirjana Efremova, Birgit Krabichler, Michael R Speicher, Johannes Zschocke, and Zlatko Trajanoski. A survey of tools for variant analysis of next-generation genome sequencing data. Brief. Bioinform., 15(2):256–278, 2014.

[11] Adam C English, William J Salerno, Oliver A Hampton, Claudia Gonzaga-Jauregui, Shruthi Ambreth, Deborah I Ritter, Christine R Beck, Caleb F Davis, Mahmoud Dahdouli, Singer Ma, et al. Assessing structural variation in a personal genometowards a human reference diploid genome. BMC Genomics, 16(1):286, 2015.

[12] John Huddleston, Mark JP Chaisson, Karyn Meltz Steinberg, Wes Warren, Kendra Hoekzema, David S. Gordon, Tina A. Graves-Lindsay, Katherine M. Munson, Zev N. Kronenberg, Laura Vives, Paul Peluso, Matthew Boitano, Chen-Shin Chin, Jonas Korlach, Richard K. Wilson, and Evan E. Eichler. Discovery and genotyping of structural variation from long-read haploid genome sequence data. Genome Res., page gr.214007.116, November 2016.

[13] Mark JP Chaisson, John Huddleston, Megan Y Dennis, Peter H Sudmant, Maika Malig, Fereydoun Hormozdiari, Francesca Antonacci, Urvashi Surti, Richard Sandstrom, Matthew Boitano, et al. Resolving the complexity of the human genome using single-molecule sequencing. Nature, 517(7536):608–611, 2015.

[14] Can Alkan, Bradley P Coe, and Evan E Eichler. Genome structural variation discovery and genotyping. Nat. Rev. Genet., 12(5):363–376, 2011.

[15] Claudia MB Carvalho and James R Lupski. Mechanisms underlying structural variant formation in genomic disorders. Nat. Rev. Genet., 17(4):224–238, 2016.

[16] Thomas Willems, Melissa Gymrek, Gareth Highnam, David Mittelman, Yaniv Erlich, 1000 Genomes Project Consortium, et al. The landscape of human STR variation. Genome Res., 24(11):1894–1904, 2014.

[17] John Huddleston and Evan E Eichler. An incomplete understanding of human genetic variation. Genetics, 202(4):1251–1254, 2016.

[18] Mark J Chaisson and Glenn Tesler. Mapping single molecule sequencing reads using basic local alignment with successive refinement (BLASR): application and theory. BMC Bioinformatics, 13(1):238, 2012.

[19] Erick W Loomis, John S Eid, Paul Peluso, Jun Yin, Luke Hickey, David Rank, Sarah McCalmon, Randi J Hagerman, Flora Tassone, and Paul J Hagerman. Sequencing the unsequenceable: expanded CGG-repeat alleles of the fragile X gene. Genome Res., 23(1):121–128, 2013.

[20] Jason Merker, Aaron M. Wenger, Tam Sneddon, Megan Grove, Daryl Waggott, Sowmi Utiramerur, Yanli Hou, Christine C. Lambert, Kevin S. Eng, Luke Hickey, Jonas Korlach, James Ford, and Euan A. Ashley. Long-read whole genome sequencing identifies causal structural variation in a Mendelian disease. bioRxiv, page 090985, December 2016.

[21] Fritz J. Sedlazeck, Philipp Rescheneder, Moritz Smolka, Han Fang, Maria Nattestad, Arndt von Haeseler, and Michael C. Schatz. Accurate detection of complex structural variations using single-molecule sequencing. Nat. Methods, Apr 2018.

[22] Heng Li. Minimap2: pairwise alignment for nucleotide sequences. Bioinformatics, 1:7, 2018.

[23] Fritz J Sedlazeck, Hayan Lee, Charlotte A Darby, and Michael C Schatz. Piercing the dark matter: bioinformatics of long-range sequencing and mapping. Nat. Rev. Genet., page 1, 2018.

[24] Adam C. English, William J. Salerno, and Jeffrey G. Reid. PB-Honey: identifying genomic variants via long-read discordance and interrupted mapping. BMC Bioinformatics, 15:180, 2014.

[25] Heng Li, Bob Handsaker, Alec Wysoker, Tim Fennell, Jue Ruan, Nils Homer, Gabor Marth, Goncalo Abecasis, and Richard Durbin. The sequence alignment/map format and SAMtools. Bioinformatics, 25(16):2078–2079, 2009.

[26] K Chidananda Gowda and Edwin Diday. Symbolic clustering using a new dissimilarity measure. Pattern Recogn., 24(6):567–578, 1991.

[27] Tobias Marschall, Ivan G Costa, Stefan Canzar, Markus Bauer, Gunnar W Klau, Alexander Schliep, and Alexander Schöonhuth. CLEVER: clique-enumerating variant finder. Bioinformatics, 28(22):2875–2882, 2012.

[28] Coen Bron and Joep Kerbosch. Algorithm 457: finding all cliques of an undirected graph. Commun. ACM, 16(9):575–577, 1973.

[29] Aric Hagberg, Pieter Swart, and Daniel S Chult. Exploring network structure, dynamics, and function using NetworkX. Technical report, Los Alamos National Lab.(LANL), Los Alamos, NM (United States), 2008.

[30] Christoph Bartenhagen and Martin Dugas. Rsvsim: an R/Bioconductor package for the simulation of structural variations. Bioinformatics, page btt198, 2013.

[31] Bianca K Stöcker, Johannes Köster, and Sven Rahmann. Simlord: Simulation of long read data. Bioinformatics, 32(17):2704–2706, 2016.

[32] Justin M Zook, Brad Chapman, Jason Wang, David Mittelman, Oliver Hofmann, Winston Hide, and Marc Salit. Integrating human sequence data sets provides a resource of benchmark SNP and indel genotype calls. Nat. Biotechnol., 32(3):246–251, 2014.

[33] Hemang Parikh, Marghoob Mohiyuddin, Hugo YK Lam, Hariharan Iyer, Desu Chen, Mark Pratt, Gabor Bartha, Noah Spies, Wolfgang Losert, Justin M Zook, et al. svclassify: a method to establish benchmark structural variant calls. BMC Genomics, 17(1):64, 2016.

[34] Miten Jain, Sergey Koren, Karen H Miga, Josh Quick, Arthur C Rand, Thomas A Sasani, John R Tyson, Andrew D Beggs, Alexander T Dilthey, Ian T Fiddes, et al. Nanopore sequencing and assembly of a human genome with ultra-long reads. Nature biotechnology, 36(4):338, 2018.

[35] Mark JP Chaisson, Ashley D Sanders, Xuefang Zhao, Ankit Malhotra, David Porubsky, Tobias Rausch, Eugene J Gardner, Oscar Rodriguez, Li Guo, Ryan L Collins, et al. Multi-platform discovery of haplotype-resolved structural variation in human genomes. bioRxiv, page 193144, 2018.

[36] Ryan K Dale, Brent S Pedersen, and Aaron R Quinlan. Pybedtools: a flexible python library for manipulating genomic datasets and annotations. Bioinformatics, 27(24):3423–3424, 2011.

[37] Michael G Ross, Carsten Russ, Maura Costello, Andrew Hollinger, Niall J Lennon, Ryan Hegarty, Chad Nusbaum, and David B Jaffe. Characterizing and measuring bias in sequence data. Genome biology, 14(5):R51, 2013.

[38] Miten Jain, Ian T Fiddes, Karen H Miga, Hugh E Olsen, Benedict Paten, and Mark Akeson. Improved data analysis for the MinION nanopore sequencer. Nature methods, 12(4):351, 2015.

