## Supplementary material for "SVIM: Structural Variant Identification using Mapped Long Reads"

#### A Supplementary figures

##### A.1 General figures

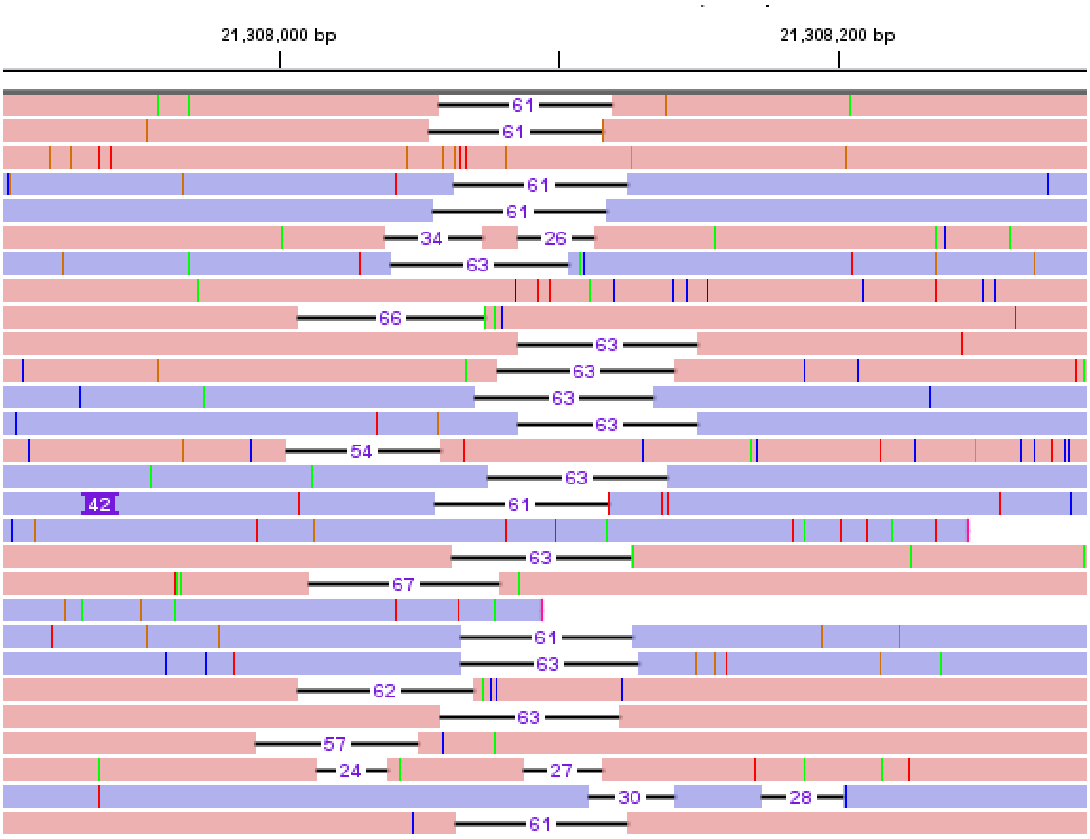


Figure S1: **The high error rate of long reads leads to a large variance in the alignment positions of deletion gaps between different reads.** This screenshot shows NGMLR alignments of PacBio reads from the NA12878 individual (viewed in the IGV genome browser). Most reads in this region of chromosome 1 (chr1:21,307,904-21,308,288) show a deletion with a length of 61-63bps but their genomic positions vary considerably. For some reads, the aligner even produced two separate gaps with a total length around 60bps to increase the alignment score.

#### A.2 Comparison of SV callers on simulated data with homozygous SVs

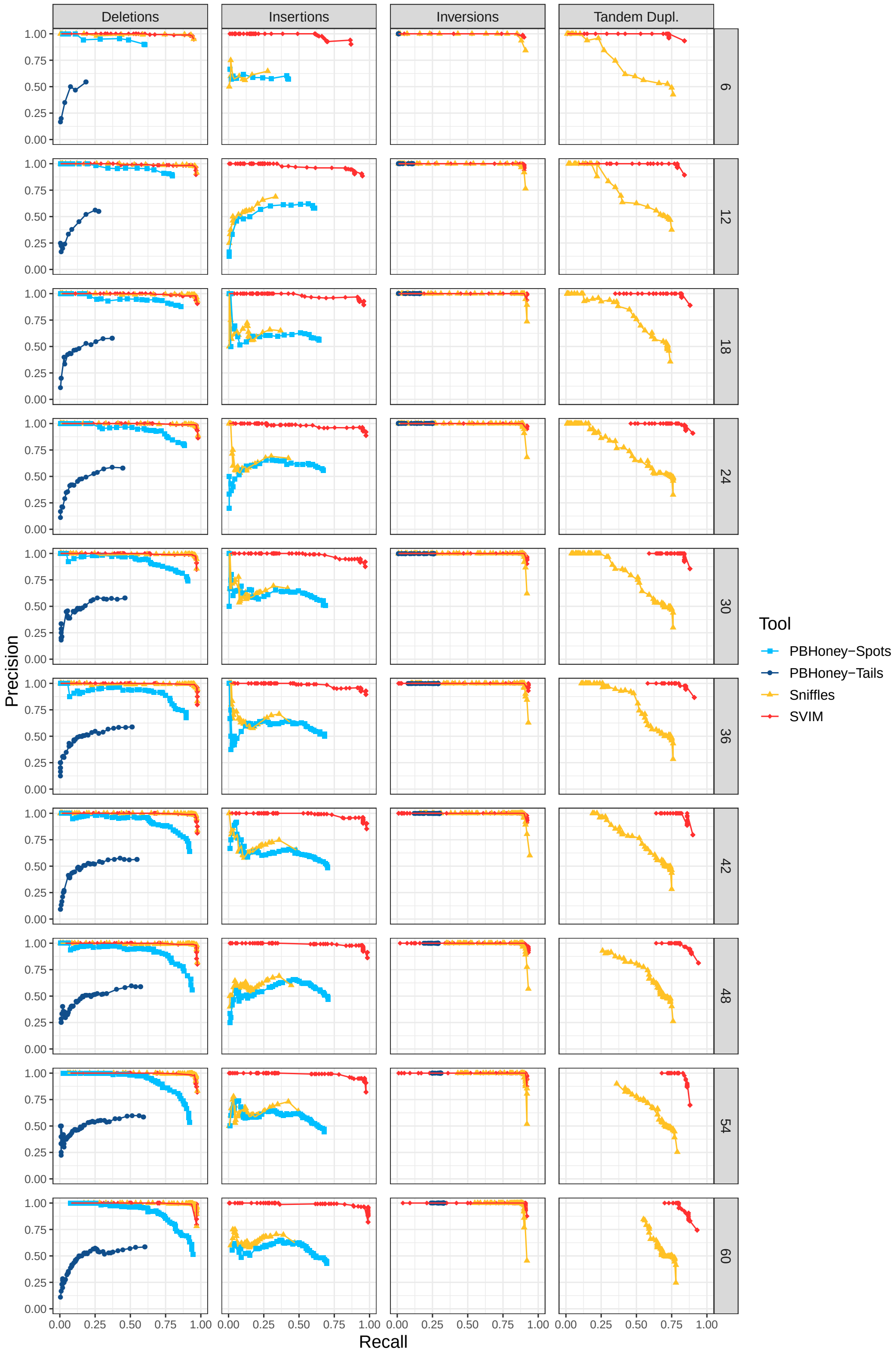


Figure S2: Comparison of SV detection performance on 10 simulated datasets with different read coverages (homozygous, alignments with NGMLR, required reciprocal overlap = 50%)

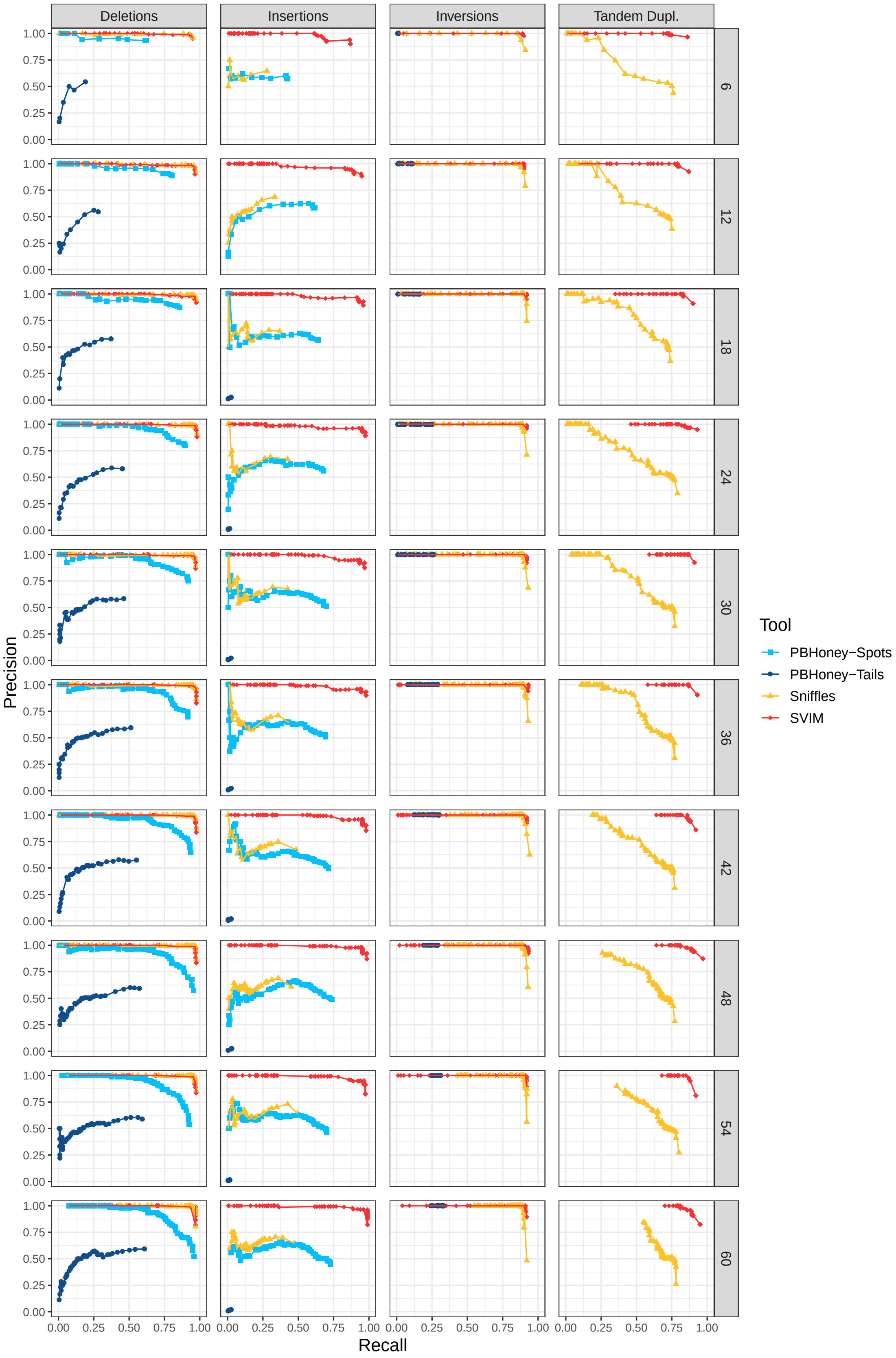


Figure S3: Comparison of SV detection performance on 10 simulated datasets with different read coverages (homozygous, alignments with NGMLR, required reciprocal overlap = 1%)

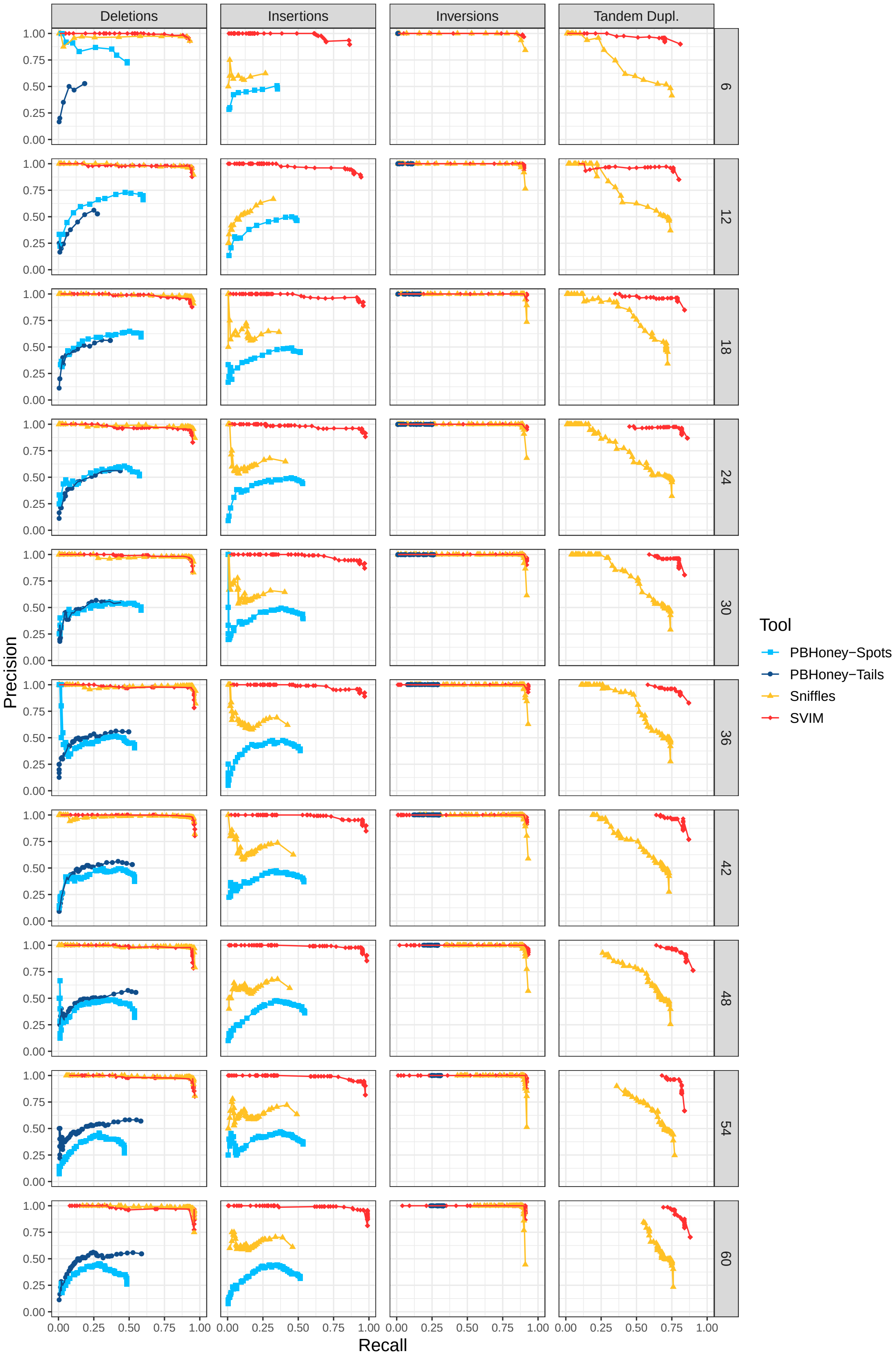


Figure S4: Comparison of SV detection performance on 10 simulated datasets with different read coverages (homozygous, alignments with NGMLR, required reciprocal overlap = 90%)

#### A.3 Comparison of SV callers on simulated data with heterozygous SVs

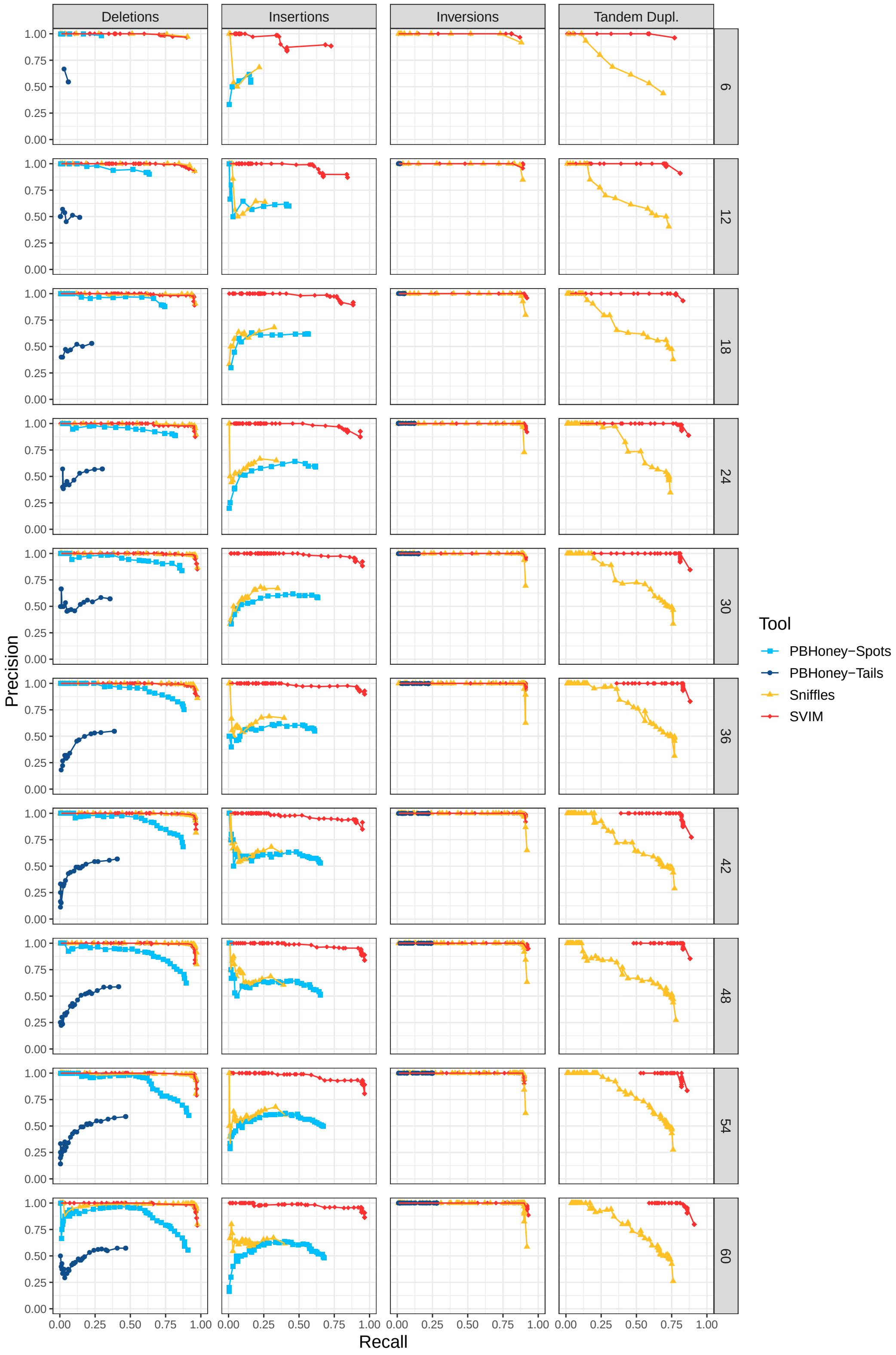


Figure S5: Comparison of SV detection performance on 10 simulated datasets with different read coverages (heterozygous, alignments with NGMLR, required reciprocal overlap = 50%)

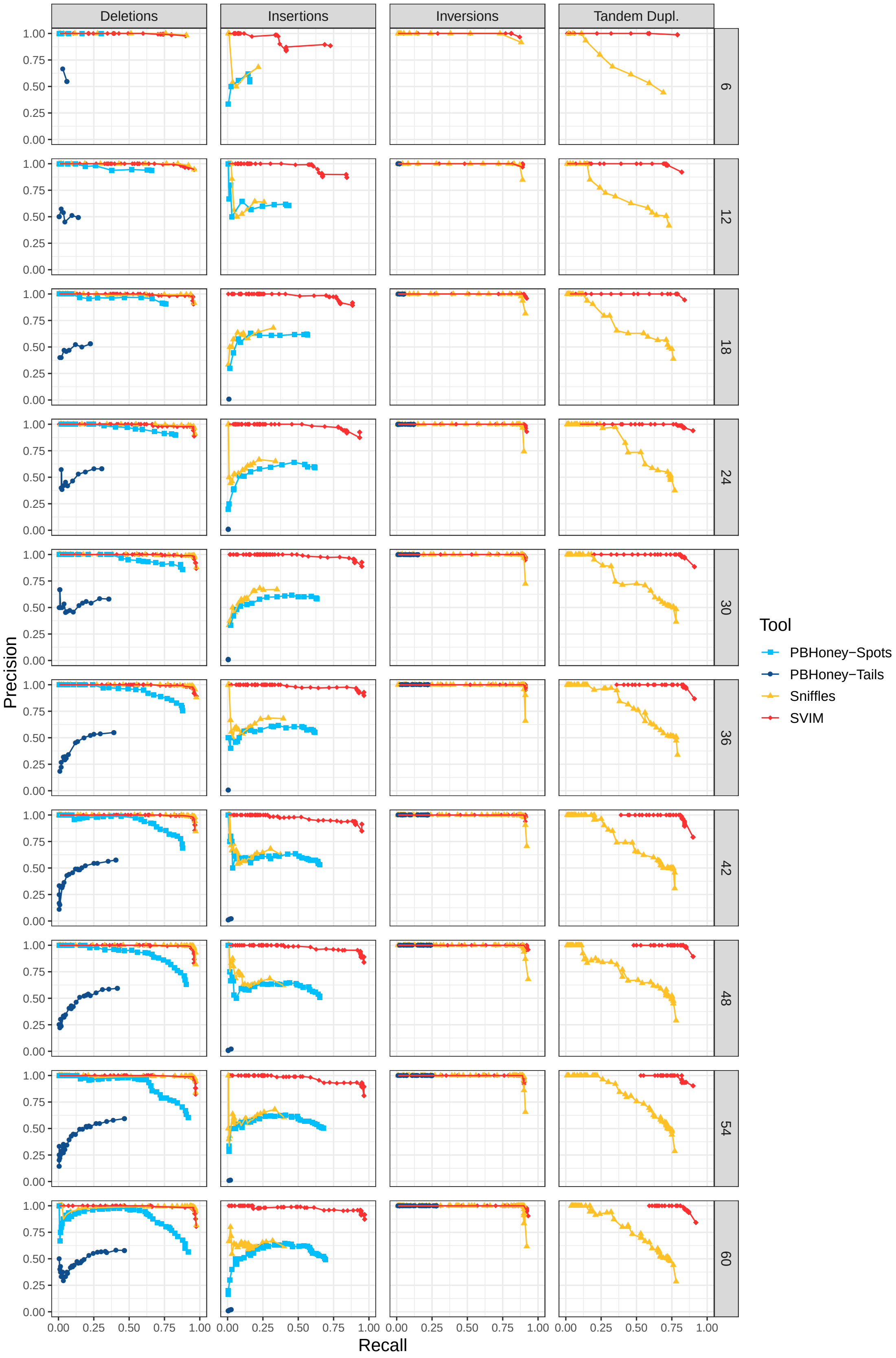


Figure S6: Comparison of SV detection performance on 10 simulated datasets with different read coverages (heterozygous, alignments with NGMLR, required reciprocal overlap = 1%)

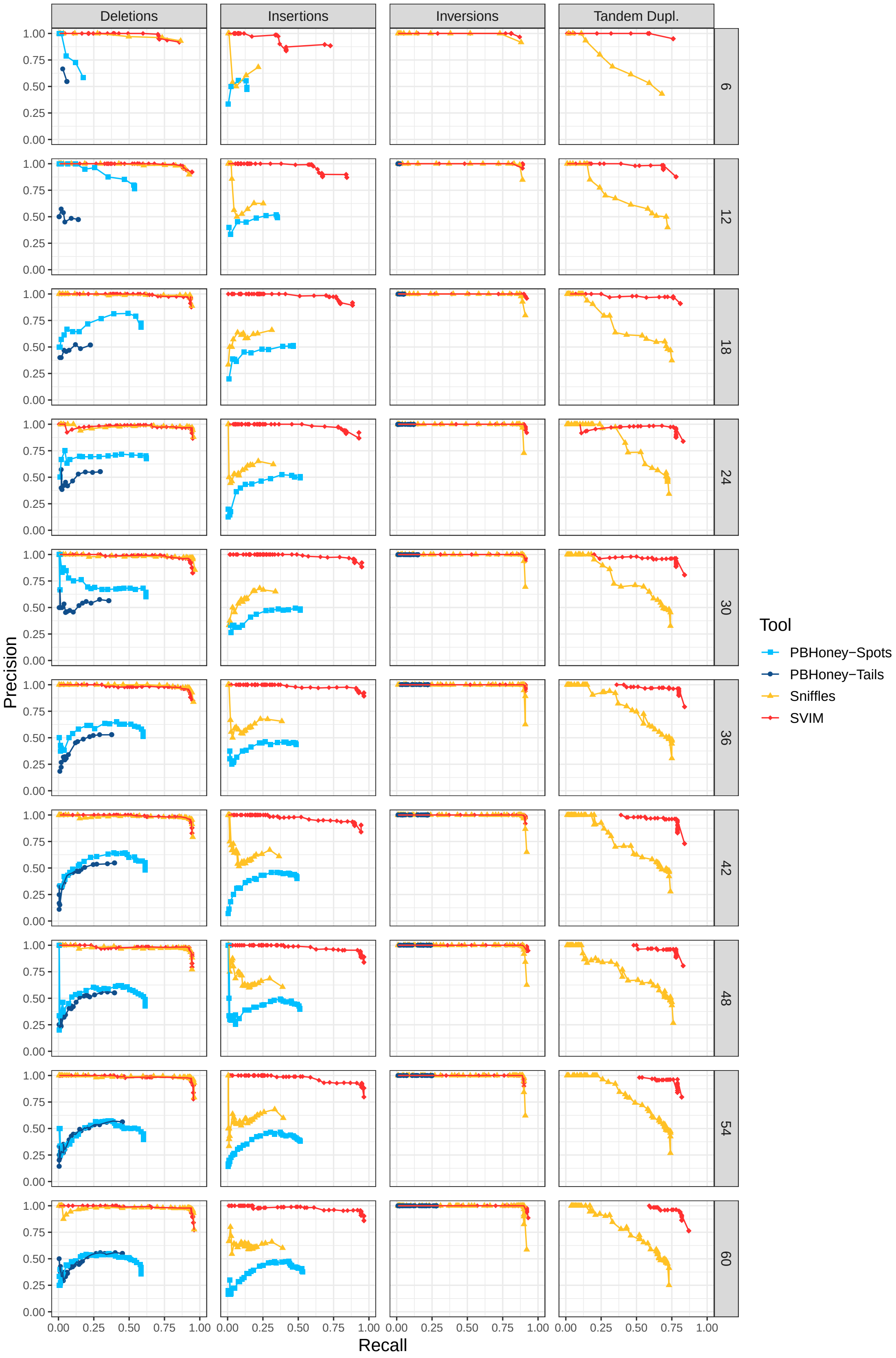


Figure S7: Comparison of SV detection performance on 10 simulated datasets with different read coverages (heterozygous, alignments with NGMLR, required reciprocal overlap = 90%)

#### A.4 Influence of read mapper on SV detection performance from simulated data

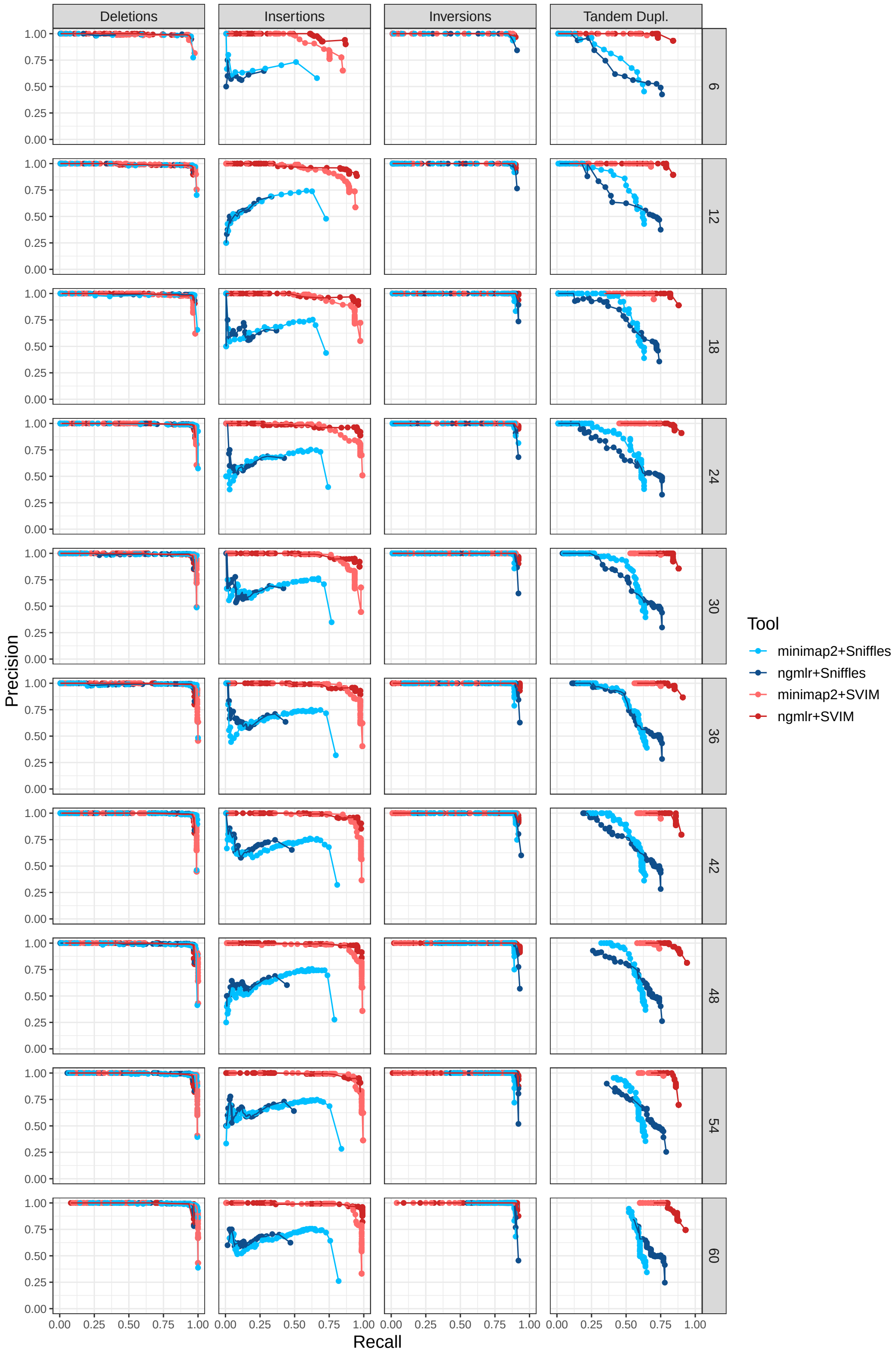


Figure S8: Evaluation of SV detection performance using alignments from two different read mappers (10 simulated datasets with different coverages, homozygous SVs, required reciprocal overlap = 50%)

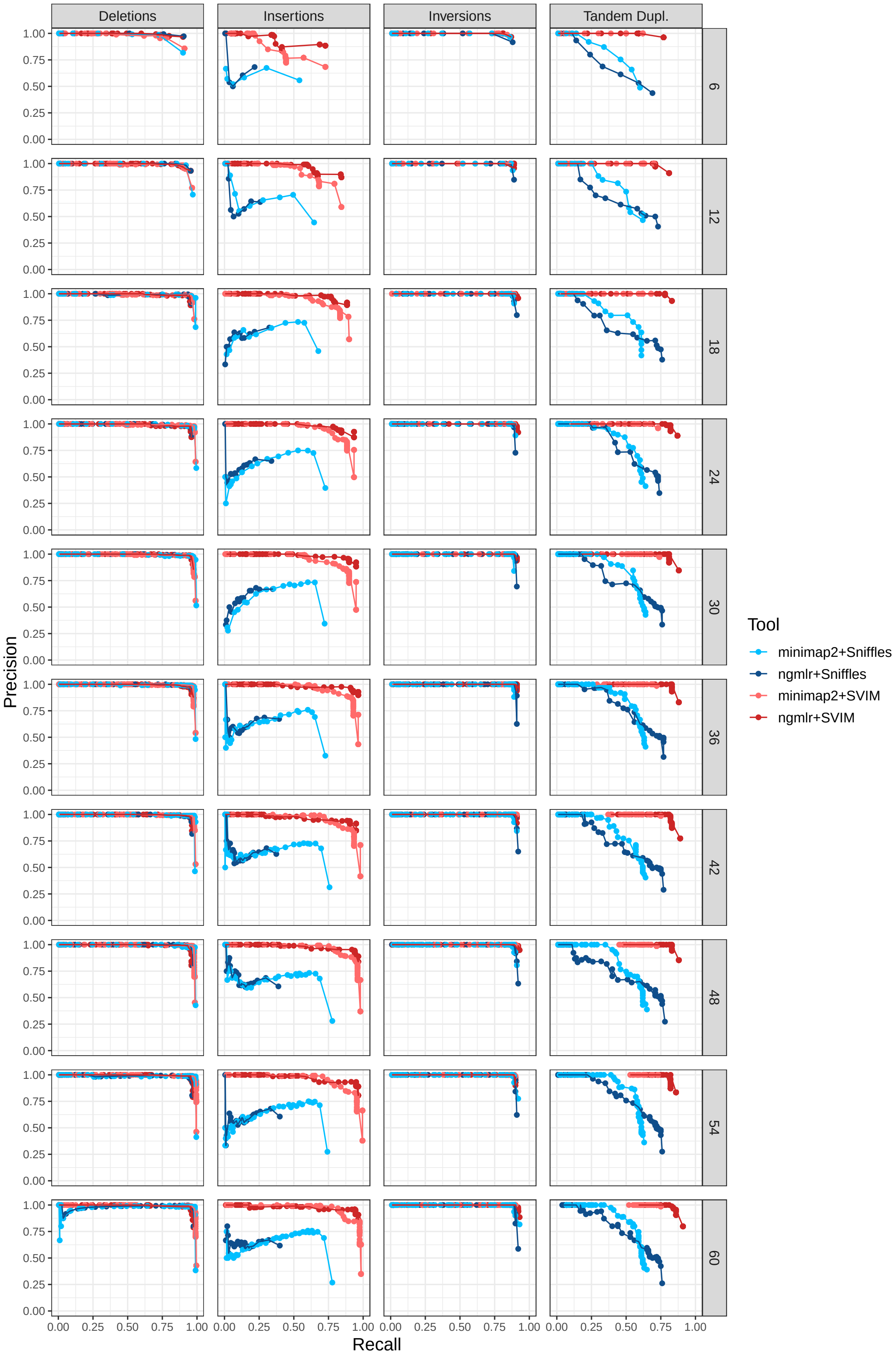


Figure S9: Evaluation of SV detection performance using alignments from two different read mappers (10 simulated datasets with different coverages, heterozygous SVs, required reciprocal overlap = 50%)

#### A.5 Comparison of SV callers on public PacBio dataset with high-confidence SVs

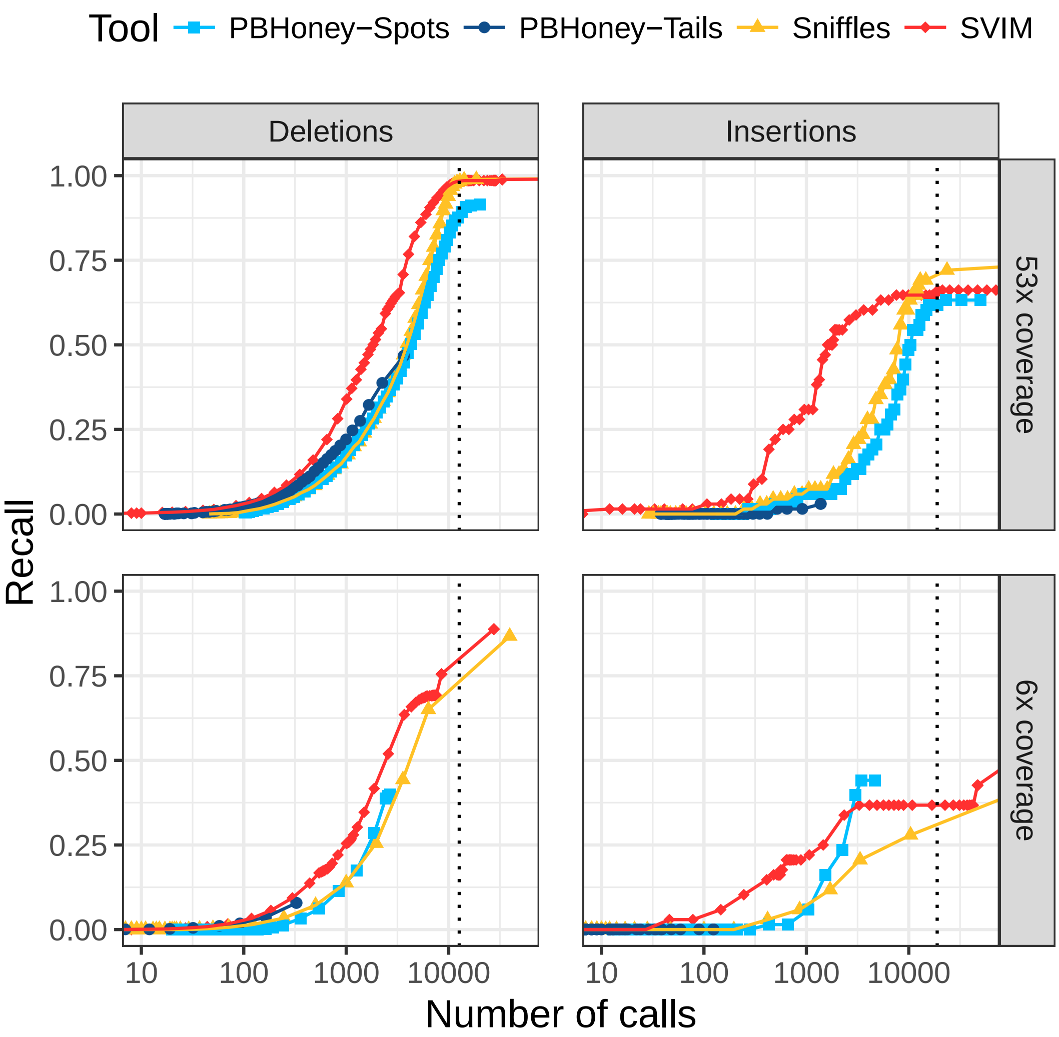


Figure S10: **Comparison of recall on a 53x coverage public PacBio dataset with 2676 high-confidence deletion and 68 insertion calls.** For each tool and different thresholds, the number of SV calls with score above the threshold (log-scale) is plotted against the recall. The upper and lower panels show performance on the full dataset and a randomly sampled 6x coverage subset of the data, respectively. SVIM reached the same recall with less calls than other tools. The vertical dotted lines denote the average number of deletions and insertions to expect in an individual as recently reported using a de-novo assembly approach (Chaisson et al., 2018). Recall was calculated using a required reciprocal overlap of 1% (deletion calls) and 1% (insertion calls), respectively, between variant calls and the gold standard variants.

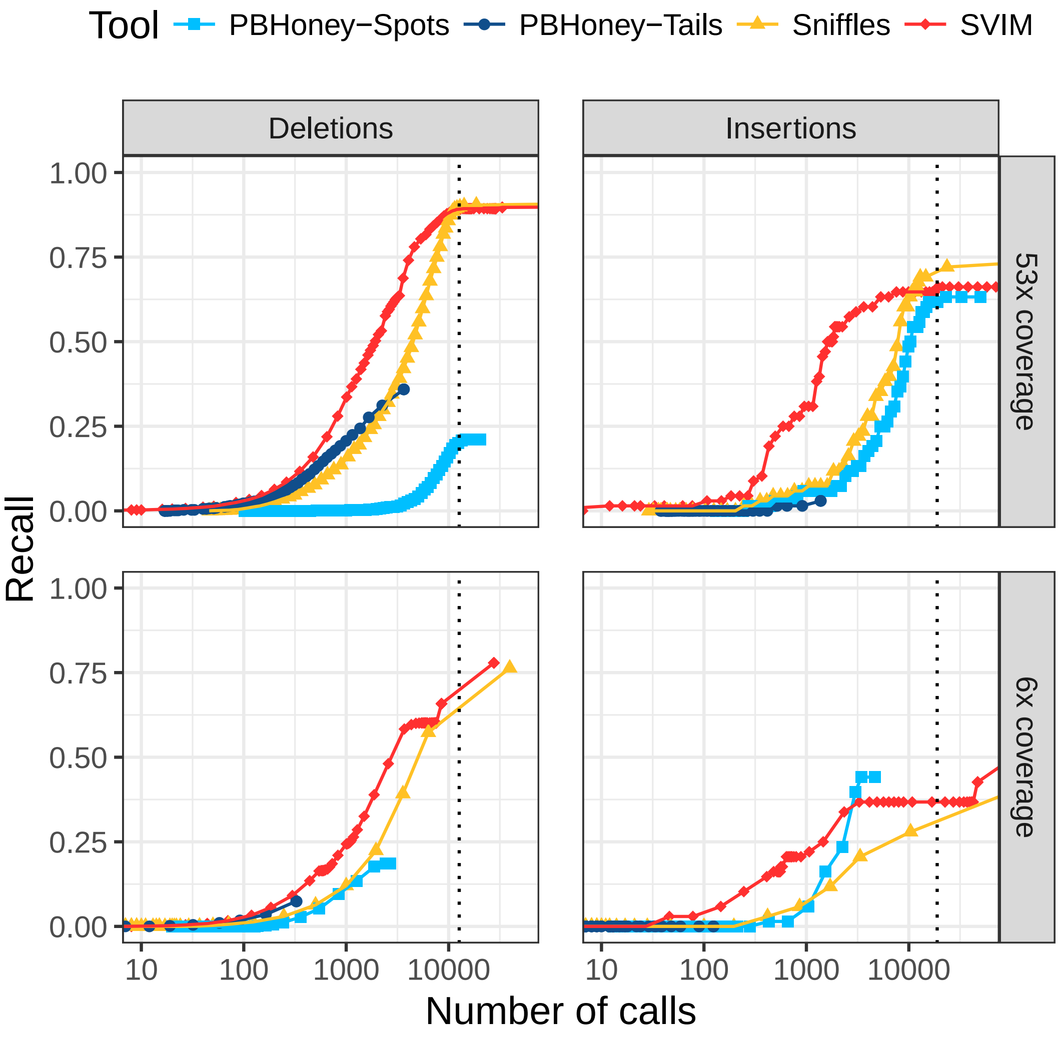


Figure S11: **Comparison of recall on a 53x coverage public PacBio dataset with 2676 high-confidence deletion and 68 insertion calls.** For each tool and different thresholds, the number of SV calls with score above the threshold (log-scale) is plotted against the recall. The upper and lower panels show performance on the full dataset and a randomly sampled 6x coverage subset of the data, respectively. SVIM reached the same recall with less calls than other tools. The vertical dotted lines denote the average number of deletions and insertions to expect in an individual as recently reported using a de-novo assembly approach (Chaisson et al., 2018). Recall was calculated using a required reciprocal overlap of 90% (deletion calls) and 1% (insertion calls), respectively, between variant calls and the gold standard variants.

##### A.6 Comparison of SV callers on PacBio reads aligned to an altered reference genome

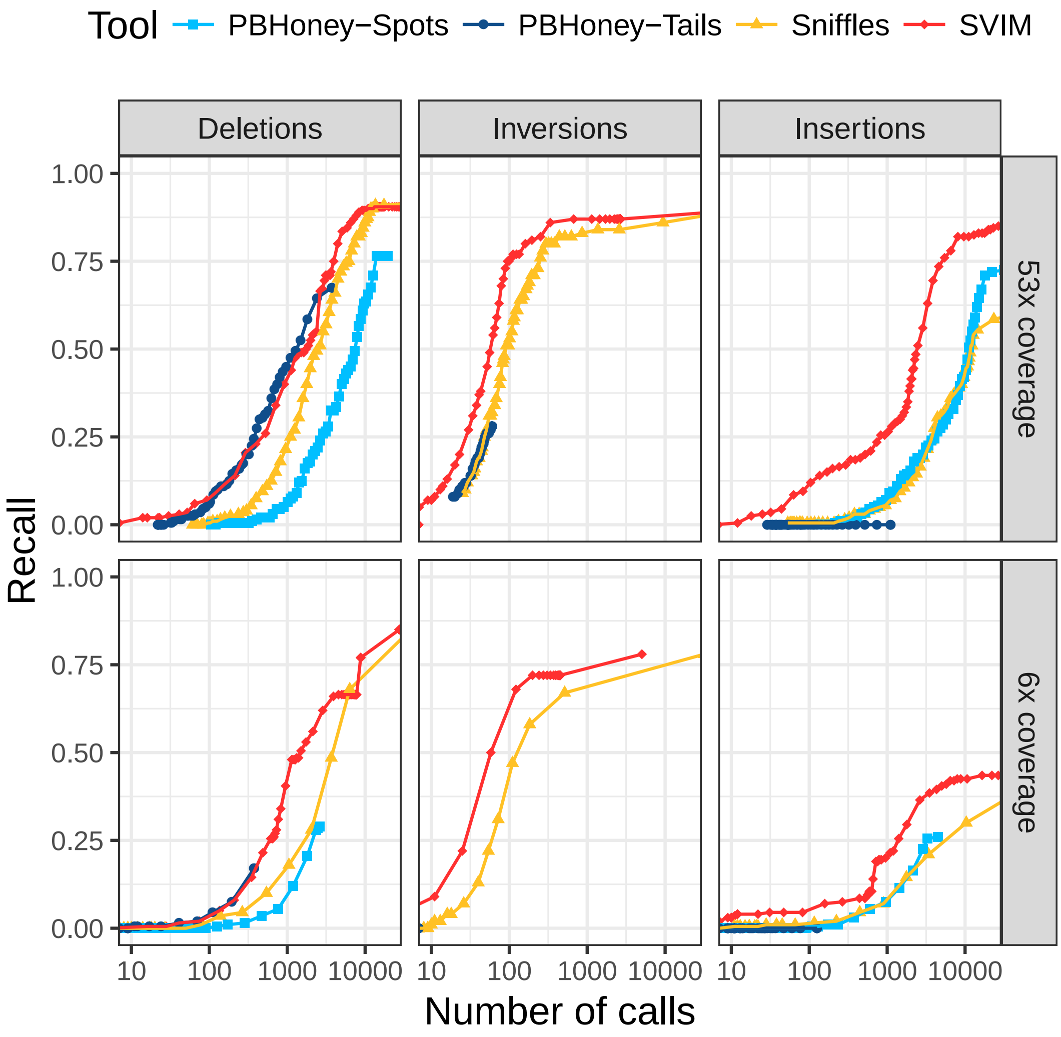


Figure S12: **Comparison of recall from NA12878 reads aligned to an altered reference genome.** For each tool and different thresholds, the number of SV calls with score above the threshold (log-scale) is plotted against the recall. The upper and lower panels show performance on the full dataset and a randomly sampled 6x coverage subset of the data, respectively. In all six panels, SVIM outperformed all the other tools and reached substantially higher recall for similar numbers of calls. The improvement was most prominent for insertions. Recall was calculated using a required reciprocal overlap of 1% between variant calls and the original implanted variants.

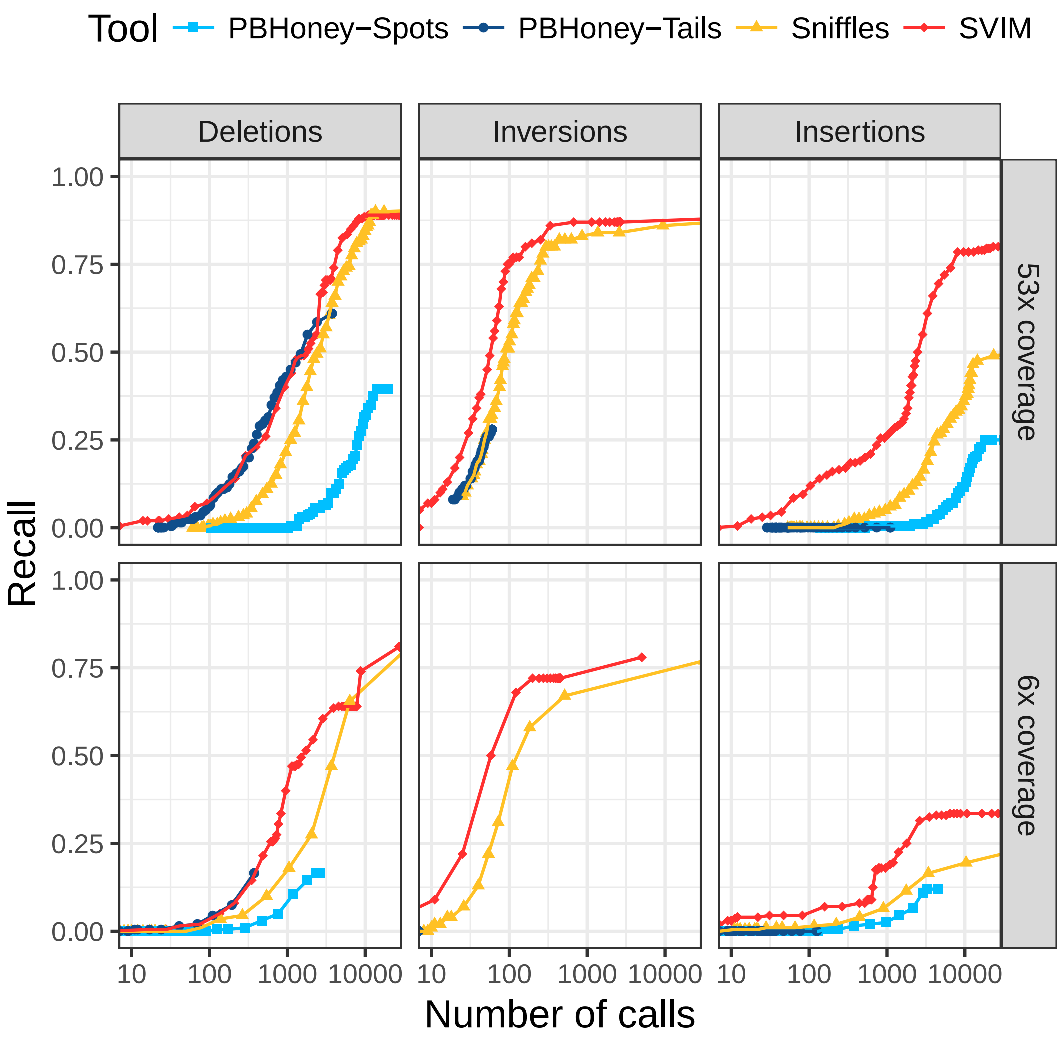


Figure S13: **Comparison of recall from NA12878 reads aligned to an altered reference genome.** For each tool and different thresholds, the number of SV calls with score above the threshold (log-scale) is plotted against the recall. The upper and lower panels show performance on the full dataset and a randomly sampled 6x coverage subset of the data, respectively. In all six panels, SVIM outperformed all the other tools and reached substantially higher recall for similar numbers of calls. The improvement was most prominent for insertions. Recall was calculated using a required reciprocal overlap of 90% between variant calls and the original implanted variants.

#### A.7 Comparison of PacBio and Nanopore callsets

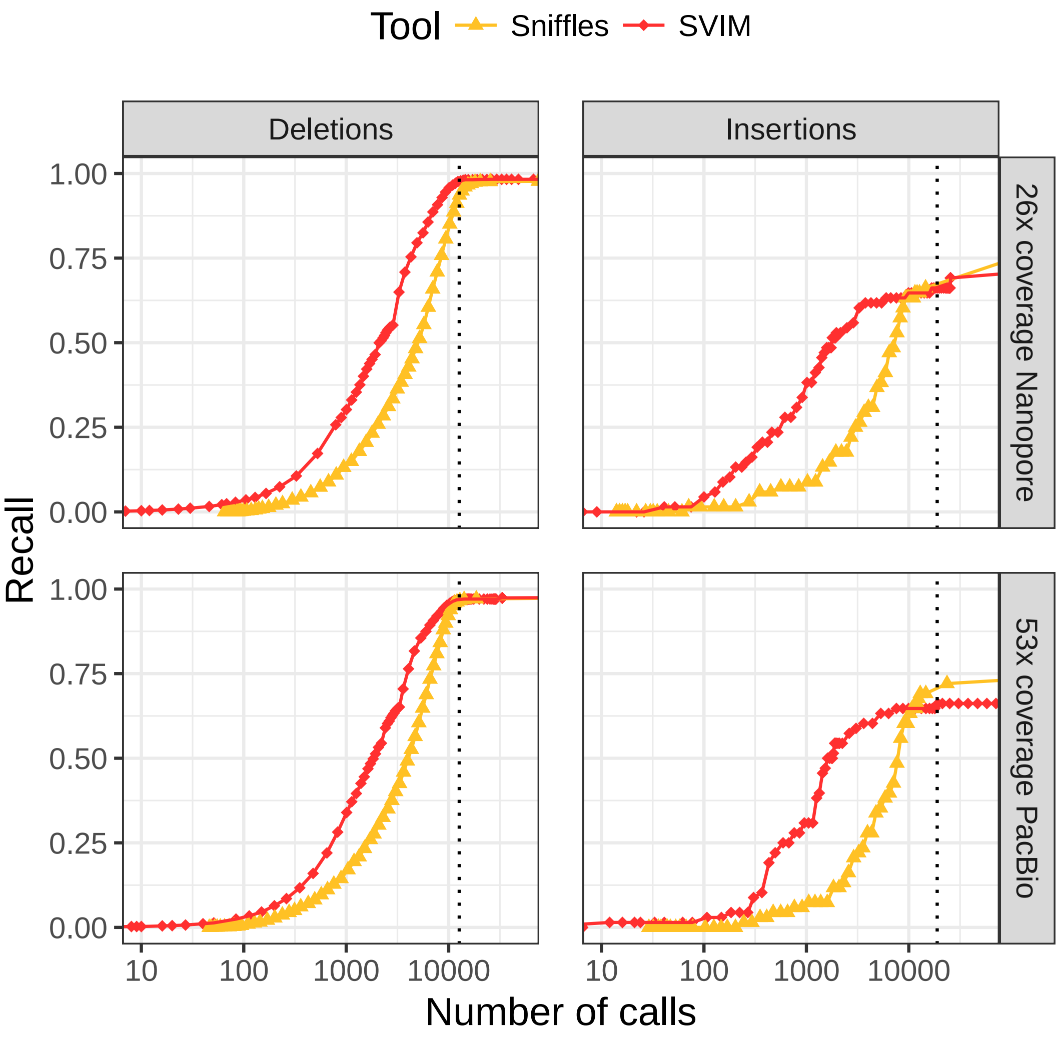


Figure S14: **Comparison of recall on a 26x coverage public Nanopore and a 53x coverage public PacBio dataset with 2676 high-confidence deletion and 68 insertion calls.** For *Sniffles* and SVIM and different thresholds, the number of SV calls with score above the threshold (log-scale) is plotted against the recall. The upper and lower panels show performance on the Nanopore and the PacBio data, respectively. For both datasets, SVIM reached the same recall with less calls than *Sniffles*. The vertical dotted lines denote the average number of deletions and insertions to expect in an individual as recently reported using a de-novo assembly approach (Chaisson et al., 2018). Recall was calculated using a required reciprocal overlap of 50% (deletion calls) and 1% (insertion calls), respectively, between variant calls and the gold standard variants.

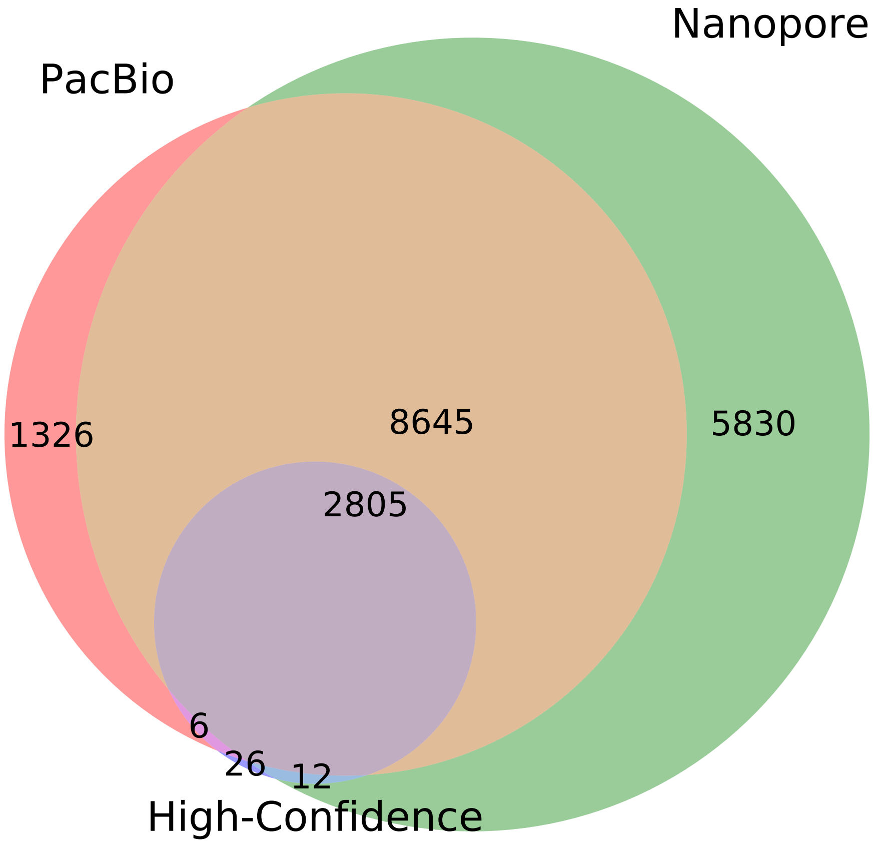


Figure S15: **Venn diagram of three deletion callsets for NA12878.** SVIM callsets on PacBio and Nanopore data as well as a high-confidence deletion callset (Parikh et al., 2016) are compared.

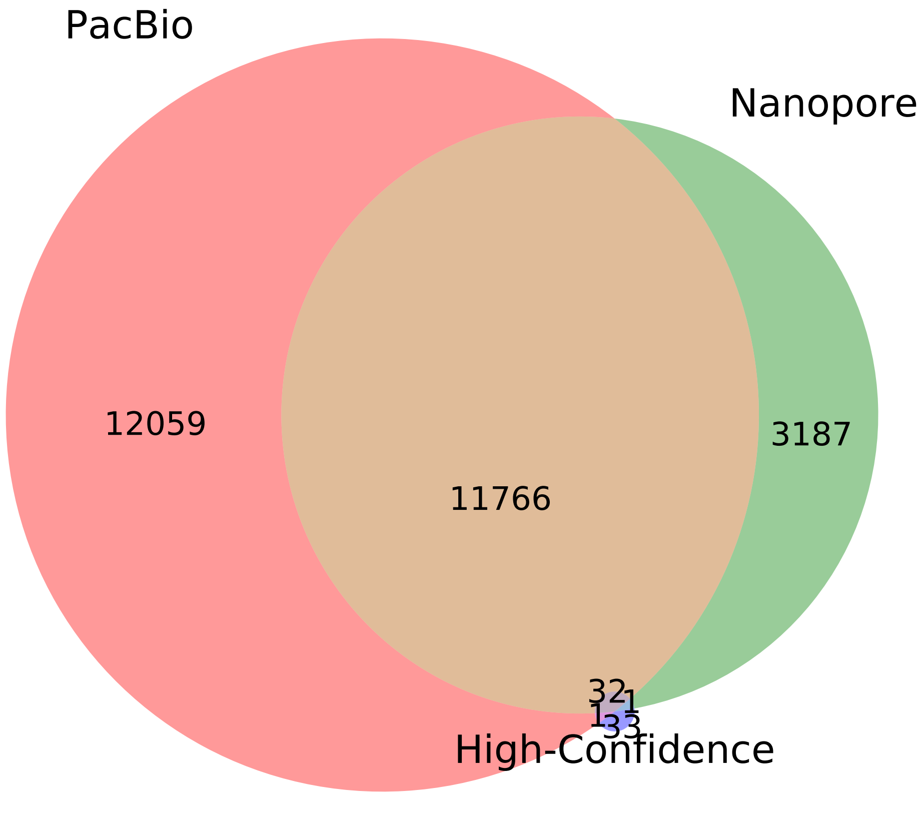


Figure S16: **Venn diagram of three insertion callsets for NA12878.** SVIM callsets on PacBio and Nanopore data as well as a high-confidence insertion callset (Parikh et al., 2016) are compared.

### B Supplementary data

Under https://www.molgen.mpg.de/~svim/svim_evaluation.zip, we provide a compressed directory with the data used for the evaluations in the manuscript. For each dataset, it contains SV call sets of the four compared tools as well as gold standard SV call sets for the different SV classes.

### List of Figures

S1 **The high error rate of long reads leads to a large variance in the alignment positions of deletion gaps between different reads.** This screenshot shows NGMLR alignments of PacBio reads from the NA12878 individual (viewed in the IGV genome browser). Most reads in this region of chromosome 1 (chr1:21,307,904-21,308,288) show a deletion with a length of 61-63bps but their genomic positions vary considerably. For some reads, the aligner even produced two separate gaps with a total length around 60bps to increase the alignment score. . . . . 1

S2 Comparison of SV detection performance on 10 simulated datasets with different read coverages (homozygous, alignments with NGMLR, required reciprocal overlap = 50%) . . . . . 2

S3 Comparison of SV detection performance on 10 simulated datasets with different read coverages (homozygous, alignments with NGMLR, required reciprocal overlap = 1%) . . . . . 3

S4 Comparison of SV detection performance on 10 simulated datasets with different read coverages (homozygous, alignments with NGMLR, required reciprocal overlap = 90%) . . . . . 4

S5 Comparison of SV detection performance on 10 simulated datasets with different read coverages (heterozygous, alignments with NGMLR, required reciprocal overlap = 50%) . . . . . 5

S6 Comparison of SV detection performance on 10 simulated datasets with different read coverages (heterozygous, alignments with NGMLR, required reciprocal overlap = 1%) . . . . . 6

S7 Comparison of SV detection performance on 10 simulated datasets with different read coverages (heterozygous, alignments with NGMLR, required reciprocal overlap = 90%) . . . . . 7

S8 Evaluation of SV detection performance using alignments from two different read mappers (10 simulated datasets with different coverages, homozygous SVs, required reciprocal overlap = 50%) . . . . . 8

S9 Evaluation of SV detection performance using alignments from two different read mappers (10 simulated datasets with different coverages, heterozygous SVs, required reciprocal overlap = 50%) . . . . . 9

S14 **Comparison of recall on a 26x coverage public Nanopore and a 53x coverage public PacBio dataset with 2676 high-confidence deletion and 68 insertion calls.** For *Sniffles* and SVIM and different thresholds, the number of SV calls with score above the threshold (log-scale) is plotted against the recall. The upper and lower panels show performance on the Nanopore and the PacBio data, respectively. For both datasets, SVIM reached the same recall with less calls than *Sniffles*. The vertical dotted lines denote the average number of deletions and insertions to expect in an individual as recently reported using a de-novo assembly approach (Chaisson et al., 2018). Recall was calculated using a required reciprocal overlap of 50% (deletion calls) and 1% (insertion calls), respectively, between variant calls and the gold standard variants. . . . . . . . . . . . . . . . . 14

S15 **Venn diagram of three deletion callsets for NA12878.** SVIM callsets on PacBio and Nanopore data as well as a high-confidence deletion callset (Parikh et al., 2016) are compared. . . . . . . . . . . . . . . . . . . . . . . . . . . . . 15

S16 **Venn diagram of three insertion callsets for NA12878.** SVIM callsets on PacBio and Nanopore data as well as a high-confidence insertion callset (Parikh et al., 2016) are compared. . . . . . . . . . . . . . . . . . . . . . . . . . . . . 16
